# A unified theory of context-conditioned efficient and predictive coding

**DOI:** 10.1101/2025.02.24.639817

**Authors:** Gaia Tavoni

## Abstract

Sensory processing does not occur in isolation: what neurons represent in a given sensory modality is shaped by signals from other senses, actions, and behavioral context. This context dependence raises a fundamental question for theories of neural coding: how can circuits efficiently encode their local input while using information available elsewhere in the brain? Here we develop a unified theory of efficient and predictive coding that shows how multimodal contextual information can optimize representations within a local sensory circuit. We demonstrate analytically that the efficient-coding solution maps onto an interpretable neural algorithm: contextual signals provide expectations about the sensory input to the local circuit, local neurons encode deviations from those expectations, and recurrent interactions whiten the residual signals. This result establishes a mathematical equivalence between context-conditioned efficient coding and predictive coding, revealing that predictive computations can emerge from efficient input compression guided by context. The resulting framework is distinct from both classical redundancy reduction within a single modality and hierarchical Bayesian inference. The theory explains and unifies diverse experimental phenomena, including cross-modal suppression of responses to predicted inputs and multimodal receptive fields across sensorimotor, audiovisual, visual–olfactory, and auditory–somatosensory circuits, while recovering classical unimodal coding effects as limiting cases. By linking coding objectives, circuit mechanisms, and experimentally observed phenomena within a single analytical framework, this work provides a principled foundation for understanding how distributed neural systems use context to shape local representations.

## I. INTRODUCTION

### A. Multimodal interactions: experimental evidence and open questions

Sensory systems have long been studied in isolation. However, converging evidence from anatomy [1, 2], neurophysiology [3–10], functional imaging [11–14], and gene expression [15] shows that signals from one sensory modality can strongly shape how stimuli are encoded across other modalities [16–19]. Multimodal interactions are thought to enhance attention to behaviorally relevant information [20], reduce perceptual noise [21], sharpen neural selectivity for preferred stimuli [9], and increase response gain – particularly when unisensory signals are weak or difficult to discriminate, a phenomenon known as “inverse effectiveness” [22]. These interactions ultimately facilitate perception [11, 23] and can arise through multiple pathways, including cortico-cortical, thalamo-cortical, and cortico-hippocampal circuits that directly or indirectly link sensory systems [1, 2, 18]. However, the specific mechanisms and computational principles underlying multimodal sensory function remain largely unknown [19, 24, 25].

Sensory processing is also profoundly shaped by movement. During movement, “efference copies” – internal representations of motor commands – are transmitted to sensory circuits, where they can either enhance [26–33] or, more commonly, suppress self-generated sensory signals across the auditory [31, 34–38], visual [39], somatosensory [40, 41], and vestibular systems [42, 43]. These mechanisms support a wide range of functions, including preventing illusory object motion during saccades [39], stabilizing gaze and posture [44], enabling vocal learning [36, 45, 46], and more generally distinguishing self-generated from external stimuli so that behaviorally relevant input is selectively processed. Beyond perception and control, recent work has further proposed efference-copy mechanisms as a foundational element in the emergence of the sense of self and conscious experience [47, 48]. Whether multisensory and sensorimotor interactions can be accounted for by common computational principles remains an open and fundamental question. Advancing neural coding theory is essential for addressing all these questions.

### B. Normative frameworks: current approaches and limitations

Over the past seven decades, three influential theoretical traditions have emerged to explain the principles of sensory coding: the efficient coding hypothesis, and two broad formulations of predictive coding.

Efficient coding posits that sensory systems maximize information transmission subject to energetic constraints, or equivalently minimize the energetic cost of encoding and transmitting a given amount of information [49–70].

Predictive coding theories propose that neural systems exploit statistical regularities to generate predictions about sensory signals. In one broad formulation, which we refer to as “retrospective predictive coding”, higher cortical areas compute estimates, or Bayesian inferences, of current or incoming stimuli, while lower sensory areas transmit the residual errors of these internal estimates [29, 30, 36, 71–80]. In this framework, prediction errors update the estimates, and the estimates, in turn, update the prediction errors through recurrent message passing across hierarchical levels, converging toward improved inferences. In a second broad formulation, which we refer to as “prospective predictive coding”, sensory systems are thought to encode information that is predictive of future stimuli or behavioral outcomes, a principle that is generally not equivalent to encoding prediction errors or unexpected information [81–87].

Despite this extensive literature, existing formulations of these theories have largely not addressed how cross-modal (sensory–sensory and sensory–motor) interactions and other sources of contextual input shape sensory codes.

### C. Context-conditioned efficient coding in multimodal canonical circuits

In this work, we develop an analytical theory of sensory coding that generalizes the efficient coding framework to circuits that process sensory inputs within a local modality while incorporating extrinsic contextual signals from other sensory modalities or motor systems. These extrinsic signals provide functional feedback that conditions processing in the local modality, irrespective of whether they are anatomically conveyed via top-down pathways. We model the local network as a recurrent circuit of reciprocally connected excitatory (E) and inhibitory (I) neurons, a widespread brain motif that serves as a fundamental processing unit across sensory systems [70, 88–94]. We refer to this type of feedback-modulated E–I network as a “multimodal canonical circuit” (Fig. 1A). For this class of circuits, we compute the connectivity configurations that achieve efficient coding of the local modality conditioned on contextual feedback in the low-noise regime.

**FIG. 1.**
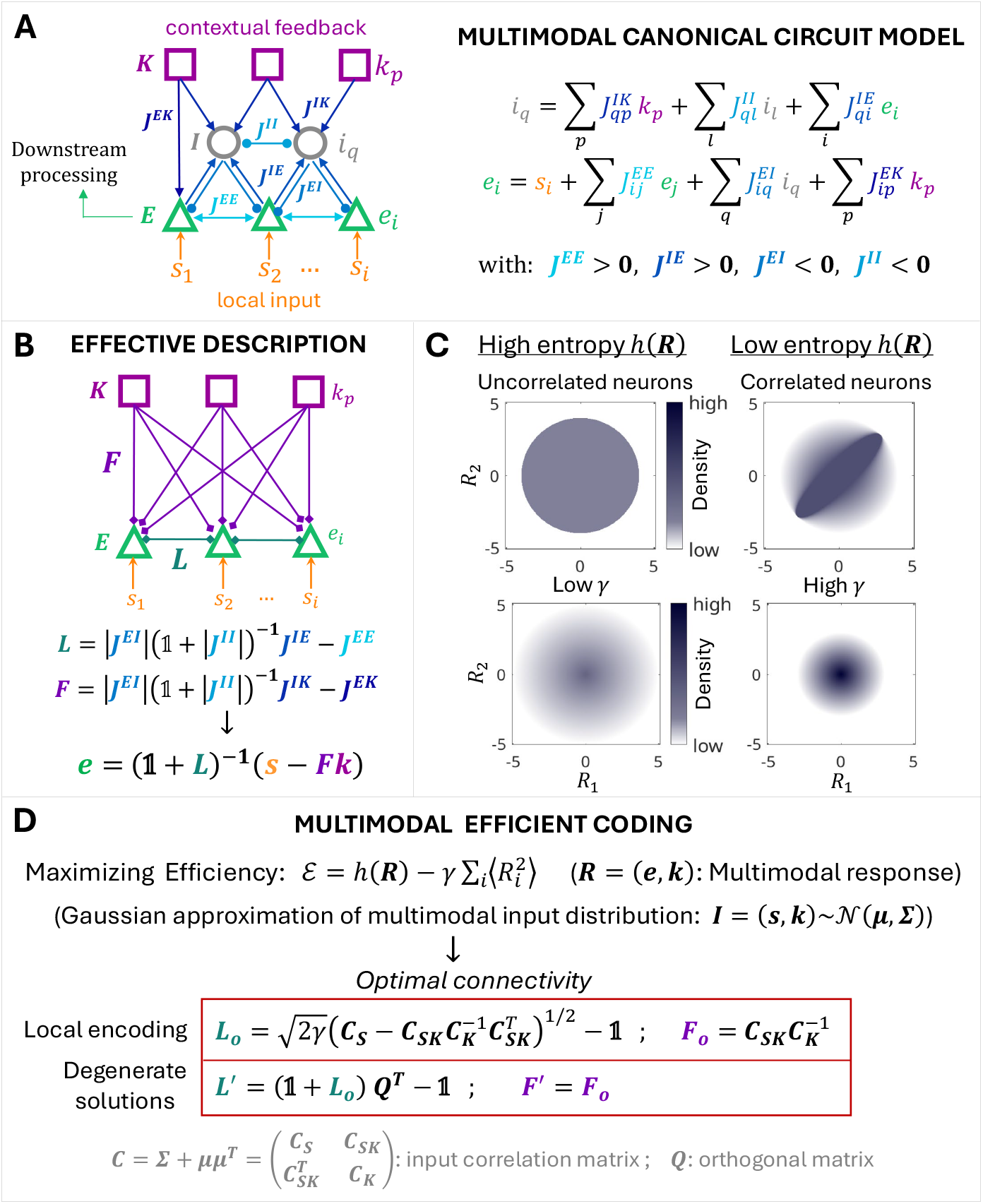
Context-conditioned efficient coding in multimodal canonical circuits. **(A)** Sketch of a multimodal canonical circuit with interconnected excitatory (E) and inhibitory (I) neurons, receiving local sensory inputs ***s*** and contextual feedback ***k*** from another population of cells (K). All inputs are linearly integrated within the circuit model. **(B)** High-level description of the circuit in (A), in which the I cells are removed, but their influence on E-cell responses is captured by two sets of effective interactions: lateral (***L***) and feedback (***F*** ) connections. These effective connections are partly mediated by I neurons and correspond to specific combinations of synaptic weights ***J*** in the detailed circuit model. **(C)** Conceptual sketch illustrating the density of multimodal representations ***R*** in neuronal response space across different scenarios. Entropy is maximized when representations are uniformly distributed across the available response space (top left). The presence of correlations between neurons increases response density in some regions while decreasing it in others, thus reducing the entropy of the representations (top right). To minimize costs, weaker responses are preferred over stronger ones, creating a gradient in response density from the center toward the periphery of response space (bottom). When maximizing Efficiency, the parameter *γ* controls the relative weighting of costs versus entropy: low values of *γ* lead to broad neuronal response ranges, expanding the available space for representing information (bottom left), whereas high values of *γ* shrink neuronal response ranges and the available space (bottom right). Increasing *γ* thus decreases costs but also reduces the achievable entropy of the representations. **(D)** Multimodal (or context-conditioned) efficient coding: mathematical formulation and optimal solution for the circuit in (A). Under a Gaussian approximation of the input distribution, the effective connectivity that maximizes Efficiency is determined by the low-order statistical structure of the multimodal inputs, represented by the ***C***_***S***_, ***C***_***K***_, and ***C***_***SK***_ blocks of the input correlation (second-moment) matrix ***C***.

### D. From the normative solution to a neural algorithm

The solution reveals a precise neural algorithm, in which each circuit component performs a precise and interpretable computation. Specifically, extrinsic feedback generates cross-modal predictions of sensory stimuli; excitatory neurons compute sensory prediction errors; and the local connectivity whitens the representation of prediction errors across the population.

This solution reveals a fundamental equivalence between context-conditioned efficient coding and retrospective predictive coding. The equivalence holds at the level of the encoded signals. However, the conceptual interpretation and the underlying algorithm differ substantially from classical, Bayesian accounts of predictive coding. In the latter, prediction errors drive reciprocal updates of higher-level estimates through iterative message passing. In our framework, by contrast, contextual signals are externally supplied conditioning variables rather than inferred latent causes; thus, prediction errors arise from context-driven compression of sensory inputs rather than from Bayesian inference over latent states.

### E. A unified account of multisensory and sensorimotor interactions

This formulation provides a parsimonious, unified account of a broad range of multisensory and sensorimotor phenomena. These include the suppression of visual responses by learned auditory cues [14]; the suppression of auditory and somatosensory responses to movement-generated sensory inputs [31, 34–38, 80, 95]; and the emergence of selective multimodal receptive fields in inhibitory neurons, as observed in the auditory [35] and olfactory systems [15]. The theory also recovers unimodal processing as a special case and reproduces classic neurophysiological findings, including range adaptation [96–98], preferential selectivity of sensory neurons for novel stimuli [99–108], and pattern separation in population responses to familiar stimuli [102, 108–110]. Finally, the theory generates new falsifiable predictions that can guide future experiments on the principles governing interactions between the senses. Among these predictions is an “inverse effectiveness” phenomenon in which contextual feedback preferentially enhances pattern separation for less discriminable sensory representations.

### F. Summary of contributions and scope

In summary, the framework proposed here represents an initial step toward integrating multimodal effects into normative theories of sensory coding. It does not aim to exhaustively characterize all possible multimodal interactions in the brain. Rather, it introduces a principled foundation that: (1) extends efficient coding to context-conditioned circuit architectures, in which coding within a local sensory circuit is optimized given extrinsic contextual signals; (2) establishes a mathematical link between context-conditioned efficient coding and retrospective predictive coding, while providing a new conceptual and algorithmic interpretation of the latter; (3) provides a unified account of a broad class of multisensory and sensorimotor effects, while recovering unimodal processing as a special case; (4) generates new, testable hypotheses about the functions and mechanisms of multimodal coding, moving beyond the traditional view of sensory systems as independent silos; and (5) offers a generalizable framework for analyzing multimodal interactions across different contextual, noise, and neurobiological constraints.

## II. RESULTS

### A. Context-conditioned efficient coding in multimodal canonical circuits

In this section, we derive the connectivity of multimodal canonical circuits that achieves efficient coding of the local sensory modality conditioned on contextual feedback.

#### 1. Multimodal canonical circuit model

We consider a canonical circuit architecture comprising reciprocally connected excitatory (E) and inhibitory (I) neuronal populations. Excitatory neurons receive feedforward inputs ***s*** from a local sensory modality, encode them in their responses (the neural representations ***e***), and transmit this information downstream. Inhibitory neurons receive extrinsic signals ***k*** from a third population, which we refer to as “context neurons” (K). These contextual signals can then modulate the representations ***e***.

Depending on the circuit, ***k*** can convey two distinct types of signals: (1) cross-modal sensory inputs, corresponding to concurrent signals from sensory modalities other than ***s***, consistent with known pathways linking different sensory systems, such as olfactory–visual [15], visual–auditory [14], and auditory–somatosensory [3, 4]; or (2) efference copies (also referred to as corollary discharges), corresponding to motor-related signals routed back to sensory areas that modulate stimulus processing during movement [31, 34–38, 80, 95]. Additional synapses can interconnect E ↔ E, I ↔ I, and K → E cells (Fig. 1A). In neocortical microcircuits, the E population can be identified with L2/3 and L5 pyramidal neurons and the I population with L1 GABAergic neurons and L2/3 interneurons, which are known to mediate inhibition from extrinsic feedback [32, 75, 92, 111–114]. Although microscopic details vary across brain regions and modalities, this circuit motif is ubiquitous and not restricted to the neocortex; closely related architectures are also found in structures such as the olfactory bulb.

In this class of circuits, we model the neuron responses (***e, i, k***) as linear functions of the respective inputs, which include the local inputs (***s***), the contextual feedback (***k***), and the inputs received from other cells through synaptic weights (***J*** ), Fig. 1A. All models of this kind, whether fully connected or not, share a common high-level description. In this description, the I neurons are virtually removed but their effects on the responses of other cells are conserved by introducing two sets of effective interactions (Fig. 1B): ***L***, representing the effective lateral connectivity between E cells, and ***F***, representing the effective feedback connectivity from K to E cells. In this model, the responses of E cells are given by:

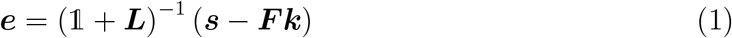

We aim to determine the effective connectivity that ensures maximally efficient coding of sensory inputs under motor or cross-modal sensory contexts.

#### 2. Multimodal (or context-conditioned) efficient coding objective

In the efficient coding framework, the optimality of a neural code is quantified by an Efficiency functional *E*, defined as the amount of information conveyed by neural responses minus the energetic cost required to represent and transmit that information. Classical formulations of efficient coding typically consider unimodal inputs and responses [51–63, 65, 66]. Here, we extend this framework to multimodal settings, in which both inputs and responses consist of a local sensory component and a contextual component. Specifically, we define the multimodal inputs and the multimodal responses as:

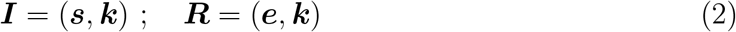

The contextual signal ***k*** thus plays a dual role: it is both a component of the organism’s overall response to the environment and an extrinsic input to the canonical circuit, which contextually modulates representations in the local modality.

Formally, the information term in *E* is quantified by the mutual information between inputs and responses, defined as the difference between the entropy of the responses and the conditional entropy of the responses given the inputs, *I*(***I***; ***R***) = *h*(***R***) − *h*(***R*** | ***I***). We consider a regime in which noise is negligible but responses are read out with finite precision (Δ). In this regime, the mutual information differs from the response entropy only by an additive constant independent of the encoding, *I*(***I***; ***R***_Δ_) = *h*(***R***) + const. We quantify energetic costs as the sum of the mean squared response amplitudes 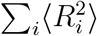 across cells, where the mean is taken over the input distribution. This constraint limits the dynamic range of neuronal responses. Dropping constant terms, we can thus define the multimodal (or context-conditioned) coding Efficiency as:

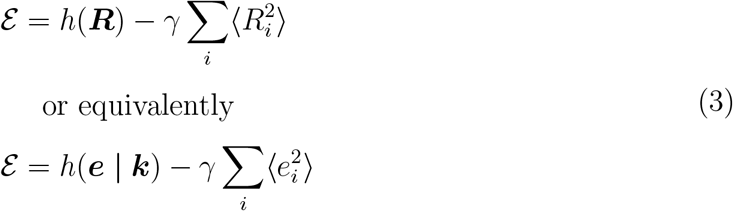

The two expressions differ only by a constant that does not affect the optimization (see Methods).

Maximizing *E* means maximizing the spread of representations (entropy) in neural response space while constraining energetic cost (Fig. 1C). The parameter *γ* is a constant that balances entropy against costs: higher values of *γ* place greater emphasis on minimizing costs, thus reducing neural response magnitudes (Fig. 1C). The second formula expresses multimodal Efficiency in terms of the components of ***R***. Since the contextual component ***k*** is extrinsic and its statistics are fixed with respect to the circuit parameters (***L*** and ***F*** ), maximizing multimodal Efficiency reduces to maximizing the conditional entropy of the local responses given the context, subject to a cost constraint on local activity. We note that a similar equivalence holds for arbitrary noise conditions, i.e., 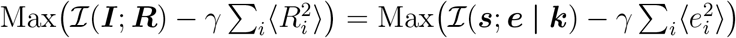 (see Methods for details).

#### 3. Context-conditioned efficient coding solution

By approximating the input distribution as a multivariate Gaussian, we can express *E* analytically as a function of the circuit connectivity and the low-order statistical structure of the multimodal inputs. Maximizing *E* yields a class of degenerate solutions with equal multimodal Efficiency. Within this class, we identify the most local encoding, defined as the solution that minimizes the mean squared distance between the inputs ***I*** and the responses ***R***. This encoding represents the minimal transformation of the inputs required to maximize multimodal Efficiency. All the other solutions differ from the most local encoding by simple rotations or reflections (Methods). Since these transformations preserve the geometry of the representation space, we can focus on the most local encoding without loss of generality: all our conclusions remain valid for the entire degenerate class. The connectivity that yields the maximally efficient and most local encoding of inputs {***s***} in the presence of contexts {***k***} is given by the following equations (Fig. 1D and Methods):

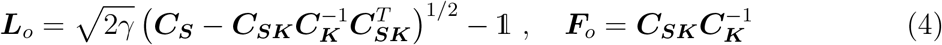

where ***C***_***S***_, ***C***_***K***_, and ***C***_***SK***_ are the blocks of the input correlation matrix ***C*** associated with the different input streams: their elements (*i, j*) represent the second moments of the input pairs (*s*_*i*_, *s*_*j*_), (*k*_*i*_, *k*_*j*_), and (*s*_*i*_, *k*_*j*_), respectively. Altogether, ***C*** characterizes the statistics of the multimodal inputs over a time window available for network adaptation. The timescales of synaptic plasticity processes in the brain constrain this adaptation window to a range of seconds to hours (though integration over longer windows may also be possible through structural plasticity). Overall, Eqs. (4) specify how multimodal canonical circuits should adapt to the low-order statistics of a local input stream and an extrinsic feedback stream over relevant timescales, which may vary across brain regions and contexts. This solution naturally generalizes to circuits receiving multiple extrinsic feedback streams (e.g., efference copies and cross-modal sensory feedback simultaneously). Specifically, different feedback streams can be accounted for as distinct blocks in the ***C***_***K***_ and ***C***_***SK***_ matrices.

We now demonstrate that this context-conditioned efficient coding solution is equivalent to a form of predictive coding driven by extrinsic context, unifying two major frameworks for understanding sensory codes.

### B. Efficient coding yields predictive codes under extrinsic context

Substituting the efficient coding solution, Eqs. (4), in the response model, Eq. (1) and Fig. 2A, we obtain (details in Methods):

**FIG. 2.**
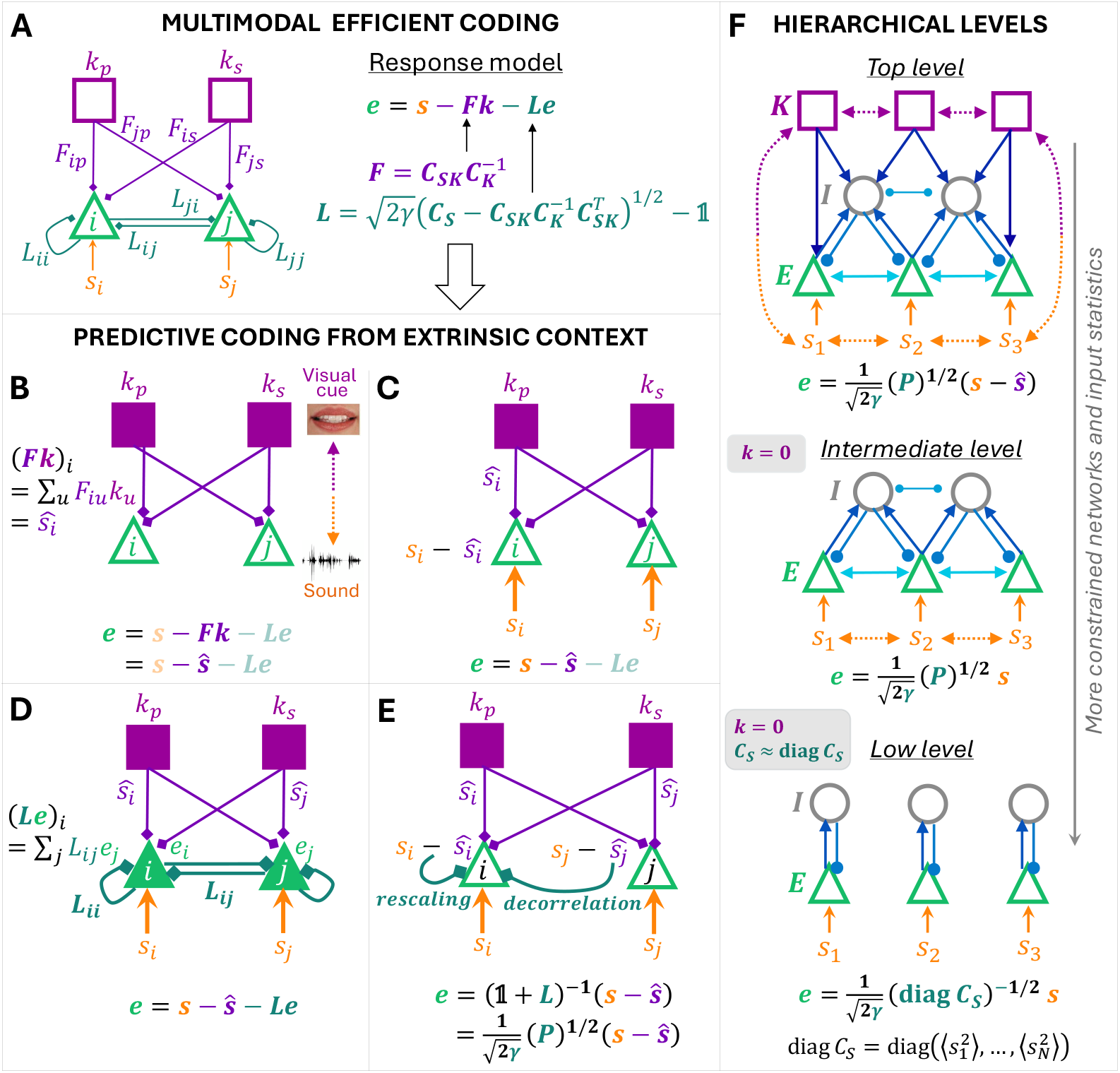
Equivalence between efficient and predictive coding under extrinsic context. **(A)** Context-conditioned efficient coding in multimodal canonical circuits. The response model, Eq. (1), describes how E cells respond to local inputs ***s***, contextual inputs ***k*** via ***F***, and lateral inputs ***e*** via ***L***. Efficient representations are achieved by substituting the optimal connectivity, Eqs. (4), into the response model. Analyzing the structure of these efficient representations reveals that efficient coding leads to predictive codes. **(B–E)**: Each component of the circuit performs an interpretable computation, establishing a precise link between efficient and predictive coding: **(B)** contextual feedback conveys optimal linear predictions {*ŝ*_*i*_} of the sensory inputs {*s*_*i*_} back to E cells. These predictions reflect learned statistical associations between local sensory inputs and extrinsic contextual signals. For example, during speech acquisition, associations form between specific phonemes (auditory inputs) and specific mouth movements (visual contextual inputs); as a result, observing those mouth movements during speech comes to predict the corresponding phonemes [116, 117]. **(C)** Extrinsic contextual predictions are subtracted from local sensory inputs in E cells, resulting in the computation of sensory prediction errors. **(D)** E cells also receive effective lateral inputs from other E cells (mediated by I neurons in the detailed circuit model). **(E)** Lateral connectivity implements the precision matrix, whitening the representation of prediction errors through rescaling and decorrelation of E-cell responses. All prediction errors are further rescaled by the constant parameter *γ*, which sets the neuronal response range. **(F)** Hierarchical levels of the theory, obtained by progressively constraining (top to bottom) network architecture and input statistics. Top level: the most general form of the theory, applicable to multimodal canonical circuits with arbitrary correlations (dotted arrows) between inputs across modalities (local and extrinsic). Intermediate level: a special case of the general theory corresponding to unimodal canonical circuits (***k*** = **0**). Low level: a special case within the intermediate-level theory corresponding to weak correlations between local inputs (***C***_***S***_ diagonally dominant). Under this condition, the effective lateral connections between E cells vanish, leaving only self-interaction terms.

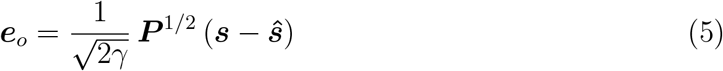

where each entry *ŝ*_*i*_ of the vector ***ŝ*** is the best prediction (minimizing mean squared error) of *s*_*i*_ that can be linearly computed from the extrinsic context ***k***, through the connectivity ***F*** . This prediction reflects previously learned statistical associations between local inputs and extrinsic contexts. In this framework, ***k*** predicts the stimulus ***s*** that is expected to be present at the same time, rather than in the future: the word ‘prediction’ is used here in the sense of ‘estimation’, consistent with retrospective predictive coding [71, 115]. ***P*** = (***C***^−1^)_[11]_ is the top-left block of the precision matrix, or inverse input-correlation matrix (hereafter, we will simply refer to ***P*** as the precision matrix). We note that ***C*** is invertible if no linear combination of inputs can perfectly predict another input in the set (***s, k***). This condition typically holds in the brain, as even a very small amount of noise in neural spiking can make activity at one site not perfectly predictable from activity at other sites.

#### 1. General interpretation of the predictive coding formula

Eq. (5) implies that E neurons encode and transmit specific linear mixtures of prediction errors. This encoding has two key properties.

First, the population representation is whitened with respect to the second-moment matrix ***C***_***ε***_ = ⟨***εε***^*T*^ ⟩ of prediction errors ***ε*** = ***s*** − ***ŝ***, since the precision matrix satisfies 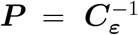 (Methods). Thus, the prediction-error mixtures conveyed by different neurons are mutually uncorrelated and have equal second moments: 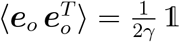.

Second, the Euclidean norm of the population response, ∥***e***_*o*_∥, encodes the magnitude of the prediction-error vector ***ε*** relative to its typical scale: 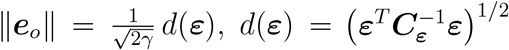. In one dimension, this reduces to a simple scale-normalized error: 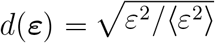. In higher dimension, *d*(***ε***) defines a Mahalanobis-like distance from the zero-error point, measured in units set by the second-moment structure of prediction errors. The equivalence between efficient and predictive coding also holds when the energetic penalty constrains response fluctuations rather than response magnitude (Appendix). In this case, the cost takes the form *γ* ⟨(*R*_*i*_ − ⟨*R*_*i*_⟩)^2^⟩ rather than 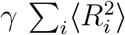 in Eq. (3). Accordingly, mixtures of prediction errors are encoded in response fluctuations rather than raw activity levels, the representation is whitened with respect to the covariance matrix of prediction errors, **Σ**_***ε***_, and the Euclidean norm of population fluctuations is 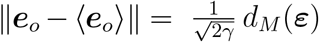. Here 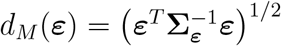 is precisely the Mahalanobis distance, that is, the distance of prediction errors from the center of their distribution (***ε*** = **0**), measured in units set by the covariance structure. This yields a direct probabilistic interpretation of the squared magnitude of response fluctuations as the negative log-likehood (or surprise) of prediction errors: 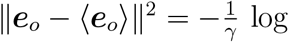 *p*(***ε***) + const.

In summary, Eq. (5) reveals a precise link between context-conditioned efficient coding and predictive coding in multimodal canonical circuits. Efficient coding naturally gives rise to predictive representations, in which each neural and network component of the canonical circuit performs a specific and interpretable computation (Fig. 2). Contextual feedback computes optimal linear predictions of sensory inputs and transmits them to E cells (Fig. 2B). Each E cell then computes a sensory prediction error by subtracting this extrinsic prediction from its local input (Fig. 2C). The local E-I network computes the precision matrix via effective lateral interactions ***L***_*o*_ among E cells. This operation rescales and integrates prediction errors across the population, yielding a whitened representation (Fig. 2D–E). Finally, the magnitude of the population response encodes the size of prediction errors relative to their typical scale. We emphasize that these predictive computations emerge directly from the efficient coding principle under extrinsic context.

#### 2. Element-wise interpretations under structured precision matrices

Under specific structural assumptions on the precision matrix, Eq. (5) can be rewritten in element-wise form, yielding distinct expressions with readily interpretable meanings. We first consider the case in which the precision matrix exhibits a dominant-mode structure, with its eigenvalues clustering into two groups: *λ*_*k*_ ≈ *λ*_1_ (the leading eigenvalue) or *λ*_*k*_ ≪ *λ*_1_. In this regime, the matrix square root simplifies to 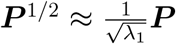. The entries of ***P*** can be expressed in terms of partial second moments *m*_*ii*·{l≠ *i*}_ and scale-normalized partial cross-moments 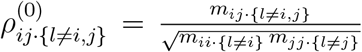, i.e., the residual moments after accounting for the linear effects of all remaining inputs *l* in (***s, k***). Each neuronal response in the vector ***e***_*o*_ can then be written as:

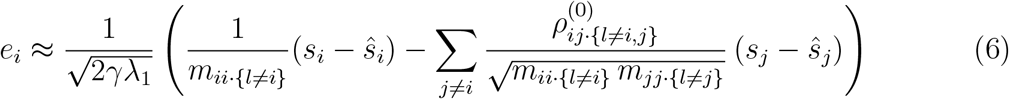

Eq. (6) shows that the E-I network scales prediction-error signals by precision-like weights derived from the inverse of their residual second moments, and combines them according to the conditional dependency structure encoded in the precision matrix. This transformation emphasizes high-precision directions while strongly suppressing low-precision ones. Prediction-error components that are conditionally redundant, as captured by partial cross-moments, are attenuated, while the representation is concentrated along the most reliable directions in input space. Finally, normalization by 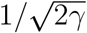 (which applies in the general case, Eq. (5)) reduces the overall response magnitude as the energetic cost of neural activity increases.

In the Methods section, we provide a second – entirely equivalent – interpretation of Eq. (5) in this low-rank regime, which emphasizes the similar roles of lateral and feedback signals in extracting redundant structure from the inputs.

We next consider the case in which the precision matrix is diagonally dominant. In this regime, Eq. (5) can be expressed in the following form:

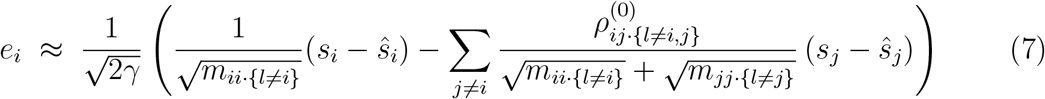

Here, each unit primarily represents its own scale-normalized prediction error, while lateral interactions introduce small subtractive corrections that suppress residual redundancy between channels. In contrast to the low-rank case, where whitening is mediated by a population-level projection onto the dominant mode, whitening here is primarily achieved at the local level.

These two regimes correspond to distinct but biologically meaningful scenarios. A dominant-mode structure in the precision matrix arises when prediction errors are strongly constrained along a small number of directions, as occurs when contextual signals accurately predict specific components of the sensory input. In such cases, residual variability is minimal along those directions, leading to dominant modes in ***P*** . In contrast, a diagonally dominant precision matrix arises when residual variability is weakly correlated across features, as may occur in highly variable environments or after substantial redundancy reduction has already taken place through early sensory processing. In this regime, prediction errors are encoded largely independently across neurons.

#### 3. Hierarchical levels of the theory

Imposing specific constraints on network properties or input statistics collapses subsets of model parameters and induces distinct hierarchical levels within the theory (Fig. 2F). Eq. (5) corresponds to the most general form of the theory (top level), which includes canonical unimodal networks as a special case (intermediate level). This intermediate level is obtained by setting all *k*_*i*_ = 0, which results in all predictions *ŝ*_*i*_ = 0. This form applies in cases where cross-modal feedback is anatomically absent or transiently non-predictive (e.g., when stimuli in two modalities are statistically independent). The intermediate level further encompasses, as a special case, canonical unimodal networks (all *k*_*i*_ = 0) with weakly correlated sensory inputs, a condition in which the off-diagonal elements ⟨*s*_*i*_*s*_*j*_⟩ of the correlation matrix ***C***_***S***_ are much smaller than the diagonal elements, 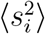. This condition corresponds to the low level of the theory. These levels do not necessarily map onto higher-to-lower anatomical regions, but instead reflect progressively more constrained network architectures and input statistics. At each level, the theory yields specific predictions appropriate to the corresponding class of networks or input statistics, as analyzed below.

### C. A unified normative account of diverse experimental findings

Across its three levels, our theory offers a unified framework for interpreting a wide range of experimental observations, including multisensory and sensorimotor interactions, and classical neurophysiological findings in individual sensory systems.

#### 1. A unified account of multisensory and sensorimotor interactions

Fig. 3 shows examples of experimentally observed multimodal effects [14, 15, 35, 37] that are reproduced by the theory at the top level (Eq. (5)) in numerical simulations of analogous experimental paradigms (see Methods).

**FIG. 3.**
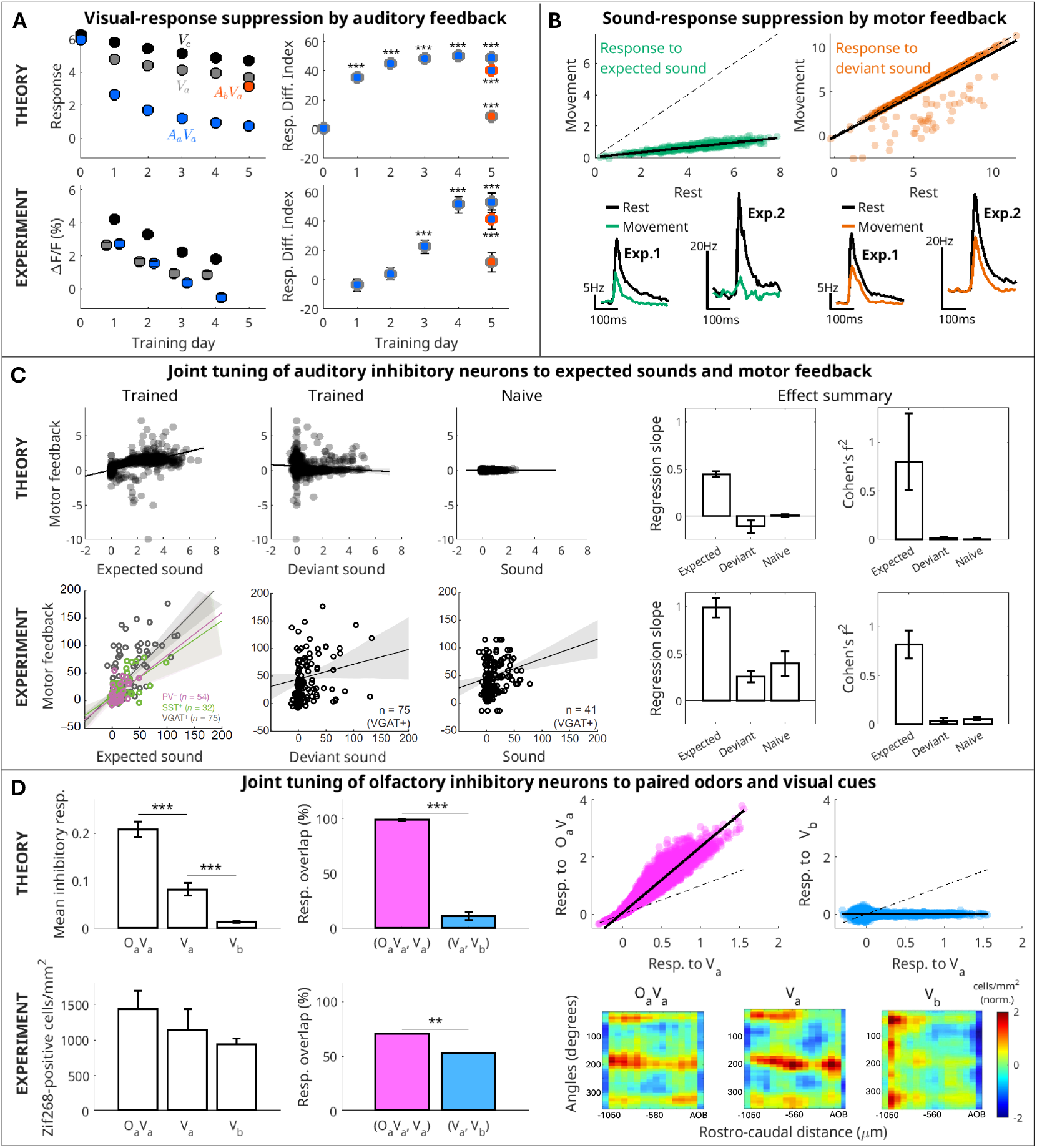
The theory provides a unified normative account of multisensory and sensorimotor neural response modulations. In each panel: top, predictions of our theory; bottom, experimental observations (reproduced with permission of Springer Nature BV, Elsevier Science & Technology Journals, and *Front. Behav. Neurosci*. from [14], [37], [35], [15]). **(A)** Audio-visual example. Left: The theory captures suppression to auditory-cued visual stimuli (*A*_*a*_*V*_*a*_) in V1 over 5 training days, stronger than for visual stimuli presented alone (*V*_*a*_, *V*_*c*_) or for an unfamiliar audiovisual pair (*A*_*b*_*V*_*a*_; Δ*F/F* data not available) [14]. Right: Response Difference Index (mean ± SE over cells and repetitions, Methods), between *V*_*a*_–*A*_*a*_*V*_*a*_ (blue circles, gray contour), *A*_*b*_*V*_*a*_–*A*_*a*_*V*_*a*_ (blue circles, orange contour), and *V*_*a*_–*A*_*b*_*V*_*a*_ (orange circles, gray contour). Here and in the following: ^∗∗∗^ *p <* 0.001, ^∗∗^ *p <* 0.01 (two-sided rank-sum test). **(B)** Audiomotor example: E responses. The theory reproduces the stronger suppression of E responses in auditory cortex to expected, compared with deviant, sounds during movement relative to rest, as in [37] (Exp. 1) and [35] (Exp. 2). **(C)** Audiomotor example: I responses. Left: The theory reproduces the stronger correlation between inhibitory responses in auditory cortex to expected sounds and motor feedback in trained animals, compared with the weaker correlation for deviant sounds or in naive animals [35]. Black lines and shaded areas: linear regressions and 95% CI. Different colors: different inhibitory neuron types. Right: Regression slopes and Cohen’s *f*^2^ (error bars: 95% CI). **(D)** Visual–olfactory example: I responses. The theory reproduces the following trends: Left: response strength (mean ± SD over repetitions) in the GC layer of the OB to *O*_*a*_*V*_*a*_, *V*_*a*_, and *V*_*b*_; Middle: significantly higher overlap of response patterns for (*O*_*a*_*V*_*a*_, *V*_*a*_) than for (*V*_*a*_, *V*_*b*_). Right: scatter plots of simulated responses (top) and experimental maps of Zif268-positive cell density (bottom).

In [14], mice were exposed to auditory-cued visual stimuli (*A*_*a*_*V*_*a*_, *A*_*b*_*V*_*b*_) that were reinforced over 5 training days, to the same visual stimuli in the absence of auditory cues (*V*_*a*_, *V*_*b*_), to a control visual stimulus (*V*_*c*_) that was never paired with an auditory cue or reinforced, and, on the last training day, to an unfamiliar audiovisual stimulus pair (*A*_*b*_*V*_*a*_). The authors found that responses in primary visual cortex (V1) to the familiar, reinforced auditory-cued visual stimuli were increasingly suppressed over the 5 training days. This suppression was mediated by long-range cross-modal connections from auditory cortex to V1. The effect was significantly stronger than in all other stimulus conditions, that is, stronger than the suppression observed (e.g., due to habituation or other factors independent of learned cross-modal feedback) when visual stimuli were presented alone (*V*_*c*_ or *V*_*a*_, *V*_*b*_ conditions) or with a novel auditory cue (*A*_*b*_*V*_*a*_ condition). Our theory closely reproduces these trends (Fig. 3A), indicating that enhanced suppression of predicted visual responses by auditory feedback after audiovisual association can follow directly from the normative principle of efficient coding. Consistent with this principle, similar inhibitory cross-modal interactions have been observed in auditory and somatosensory cortices [3, 4]. Another class of experiments concerns sensorimotor interactions. In [35], mice learned to associate a sound with their movement on a treadmill, where tone pips were delivered at a rate proportional to running speed. Similarly, in [37], mice were trained to expect a sound at a specific time during a forelimb movement to press a lever. In both studies, neuronal activity in auditory cortex was recorded after learning, during presentation of either the expected sound or a deviant sound. Responses were measured both during movement (running or lever pressing) and at rest. In both experiments, responses to the expected sound during movement were suppressed relative to rest, and this suppression was significantly stronger than that observed for the deviant sound. Furthermore, these prediction-error responses were concentrated in layers 2/3 and 5 [37], consistent with the interpretation of the E population in our model. Similar predictive suppression driven by volitional movements has been observed in the somatosensory [40, 41, 95] and vestibular systems [42, 43, 118]. Our theory reproduces both the qualitative response patterns and their magnitude (Fig. 3B), indicating that suppression of the expected sensory consequences of movement can be explained by the normative principle of efficient coding.

In [35], the authors also recorded the activity of inhibitory neurons in the auditory cortex in response to the expected and deviant sounds, as well as to stimulation of the secondary motor cortex. While responses to the deviant sound and to motor feedback were only weakly correlated, responses to the expected sound and motor feedback exhibited a strong correlation after learning the audiomotor association. In contrast, this correlation was markedly reduced in naive animals that had not been trained to expect the sound. We used a hybrid analytical-numerical approach to infer synaptic weights satisfying Dale’s law and giving rise to effective connections that approximate the normative solution. From these inferred weights, we estimated optimal I-cell responses according to the theory (see Methods). The theory reproduces the significantly stronger correlation observed in I-cell responses to the expected sound and the motor feedback (Fig. 3C and Supplementary Fig. S1). For the deviant sound, the theory predicts variable but generally weak correlations (slightly negative or statistically zero, depending on the stimulus frequency, Fig. 3C and Supplementary Fig. S1), broadly consistent with the experimental observations. These results strongly suggest that prediction-error encoding in the E population arises from the transmission of feedback predictions via the I population, consistent with our theoretical framework.

In [15], the authors reported a similar correlation in the receptive fields of granule cells (GCs, inhibitory neurons) in the olfactory bulb (OB) with respect to paired odors and visual cues. Mice were conditioned over 10 days to associate an odor (*O*_*a*_) with a visual context (*V*_*a*_). On day 11, neural activity in the GC layer of the OB was assessed by mapping the expression of the immediate early gene Zif268. Responses were measured under three conditions: following odor presentation in the same visual context (*O*_*a*_*V*_*a*_), presentation of the visual context alone (*V*_*a*_), and presentation of a different visual context (*V*_*b*_). The authors first showed that visual contexts alone were sufficient to elicit activity in the GC layer, with activation decreasing on average from *O*_*a*_*V*_*a*_ to *V*_*a*_ to *V*_*b*_. Although the differences across conditions were not reported to be statistically significant, these patterns indicate the presence of feedback to the GC layer conveying information about visual context, and suggest that feedback representing the learned context *V*_*a*_ is strengthened during learning. Our theory predicts the same qualitative trend, emphasizing differences across the three conditions (Fig. 3D, left).

Second, the activation patterns observed in [15] for *O*_*a*_*V*_*a*_ and *V*_*a*_ were significantly more similar than those observed for *V*_*a*_ and *V*_*b*_, indicating that GCs developed cross-modal tuning selective for paired odors and visual cues, analogous to the selective audiomotor tuning of inhibitory neurons in the auditory cortex [35]. Our theory reproduces this trend as well (Fig. 3D, middle and right). We note that the overlap between I-cell responses to *V*_*a*_ and *V*_*b*_ is smaller in our simulations than in the experimental data because we modeled *V*_*a*_ and *V*_*b*_ as orthogonal feedback signals. Modeling the visual feedback signals as distinct but non-orthogonal increases the overlap in I-cell responses, consistent with the experimental observations (Supplementary Fig. S1). Non-orthogonal signals are also plausible for visual contexts that share common features (e.g., overall cage shape) while differing in others (e.g., visual cues on the cage walls), as in [15].

#### 2. A unified account of classical unimodal effects

We now consider the theory’s expected behavior at the intermediate level, corresponding to canonical unimodal networks (Fig. 2F). In this simplified form, E responses are predicted to be linear mixtures of sensory inputs: 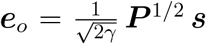. Compared to the most general expression given by Eq. (5), here the predictions ***ŝ*** are turned off. In this form, the theory accounts for habituation to repeated stimulus presentations (Fig. 4A), with neuronal tuning shifting away from familiar inputs and toward novel stimuli (Fig. 4B). Additionally, it predicts progressive pattern separation in responses to similar inputs (Fig. 4C). These effects are driven by the local E-I connectivity and align with classical neurophysiological findings across sensory systems [99–110].

**FIG. 4.**
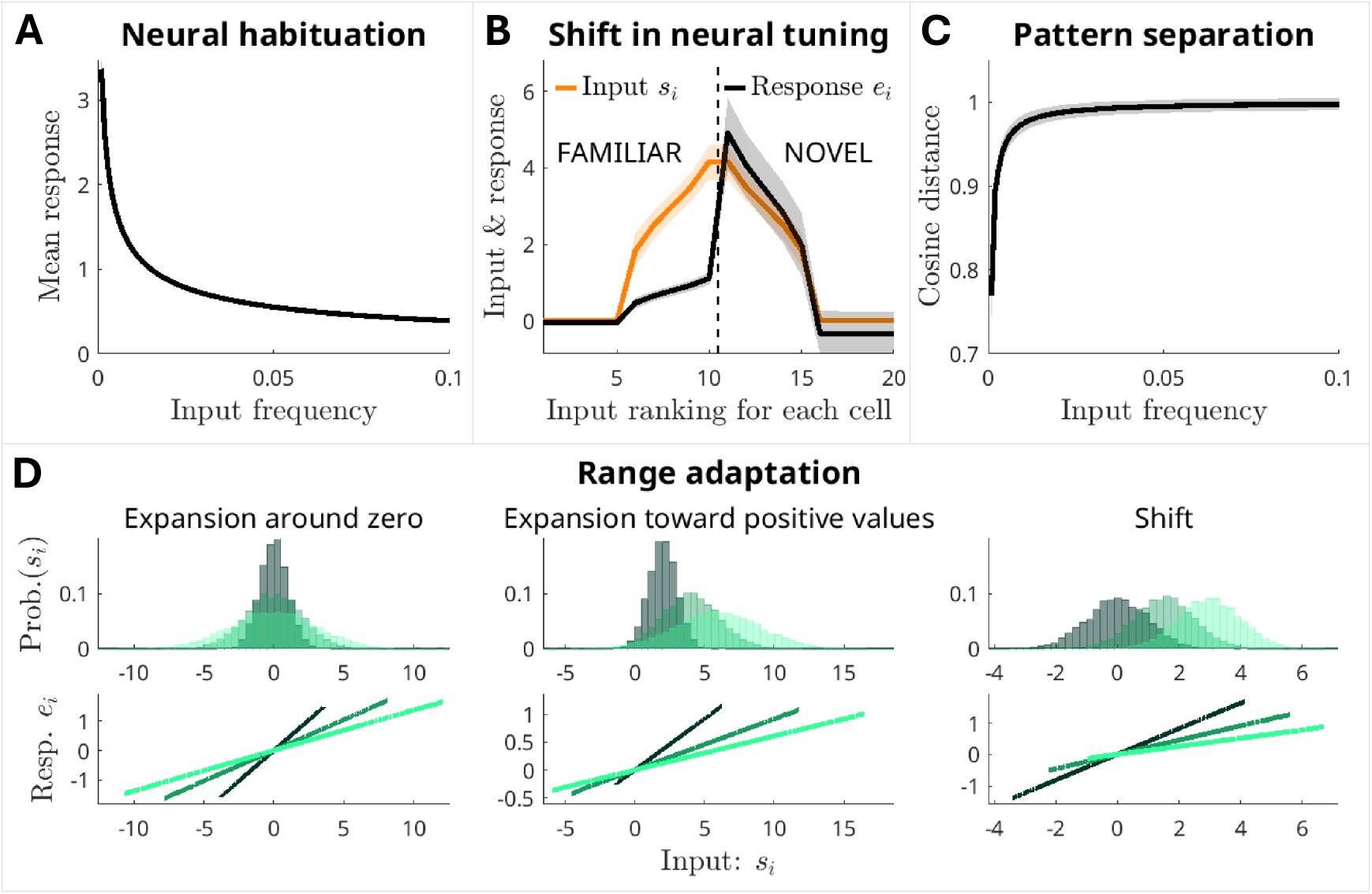
The theory provides a unified normative account of classical unimodal effects. **(A–D)** Effects predicted in the absence of cross-modal feedback. **(A)** The theory reproduces habituation effects observed across sensory systems [99–102, 106, 108]. The mean response of E cells to sensory input decreases as the input frequency increases. **(B)** The theory replicates the shift in neural tuning away from familiar inputs and toward novel stimuli observed in the olfactory, auditory, and visual systems [99–108]. The plot shows the mean and IQR of inputs (orange) and responses (black) of E cells to familiar and novel inputs, as computed from the optimized model (Methods). **(C)** The theory reproduces pattern separation effects observed in the olfactory and visual systems [102, 108–110]. The plot shows the mean and IQR of the cosine distance between E-cell responses to pairs of stimuli with 50% overlap. The stimuli are presented at increasing frequency. The cosine distance increases as the stimuli become familiar. **(D)** The predicted range adaptation effects are similar to those observed in various brain regions [96–98, 119–121]. The response of an excitatory (E) cell in the model is shown as a function of the cell’s input (bottom plots) after adaptation to the input distribution in the same color (top plots). The response gain adjusts to the typical input range.

At the lowest level (Fig. 2F), Eq. (5) reduces to 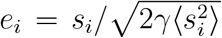, for each cell. Each input is rescaled by its root mean squared (RMS) strength, while the constant parameter *γ* regulates the typical response amplitude of all neurons. This solution embodies range adaptation, a well known phenomenon in which neurons in sensory and other cortical or midbrain areas tend to utilize their full response range to represent inputs in the current environment [96–98, 119–124]. Here, divisive normalization by the RMS input amplitude reproduces the decrease in response gain observed when the range of inputs expands around zero (Fig. 4D, left, and [97, 98]) or in the positive direction (Fig. 4D, middle, and [119, 120]). RMS normalization also promotes partial range adaptation when the input domain shifts, allowing neurons to reallocate part of their response range to represent the new input domain (Fig. 4D, right, and [96, 121]). While the low-level theory is sufficient to predict these effects, range adaptation is not exclusive to the low-level description but is carried forward and integrated with other computations at higher levels. In particular, in the general form, Eq. (5), range adaptation arises naturally from the whitening of the E responses by the square root of the precision matrix.

Our theory also generates a range of new falsifiable predictions, offering a clear path for further experimental validation.

### D. New predictions for multimodal interactions

#### 1. Context-enhanced selectivity for novel stimuli

A new prediction from Eq. (5) is that habituation and neural selectivity for novel stimuli are enhanced for greater efficiency when information is redundant across modalities. Specifically, when a context ***k*** repeatedly co-occurs with a stimulus ***s***, the redundant portion of the stimulus explained by ***k*** is subtracted from the ***e*** responses, leading to enhanced adaptation (Fig. 5A). By a similar mechanism, familiarization with different stimuli, each presented in a distinct context, sharpens neural selectivity for novel stimuli (Fig. 5B). These effects increase as the statistical association between each stimulus and its corresponding context becomes stronger (Fig. 5AB, compare ‘weak’ and ‘strong’ context). Here we refer only to the strength of the statistical association, not to the number of activated context cells or their activation strength. The overall strength of the feedback signal is determined by all these factors together. For numerical reasons, each context in Fig. 5B was simulated by the activation of a single context unit; thus, the relative strength of the feedback compared to the analyses of Fig. 3 was weaker. This explains the relatively weaker feedback-induced suppression of expected stimuli compared to the suppression shown in Fig. 3B. We also note that Fig. 5B shows the theoretical prediction for a different stimulation paradigm, in which each stimulus is associated with a different context, whereas in Fig. 3B the context was fixed (represented by the same learned movement) and only the familiarity of the stimulus was varied.

**FIG. 5.**
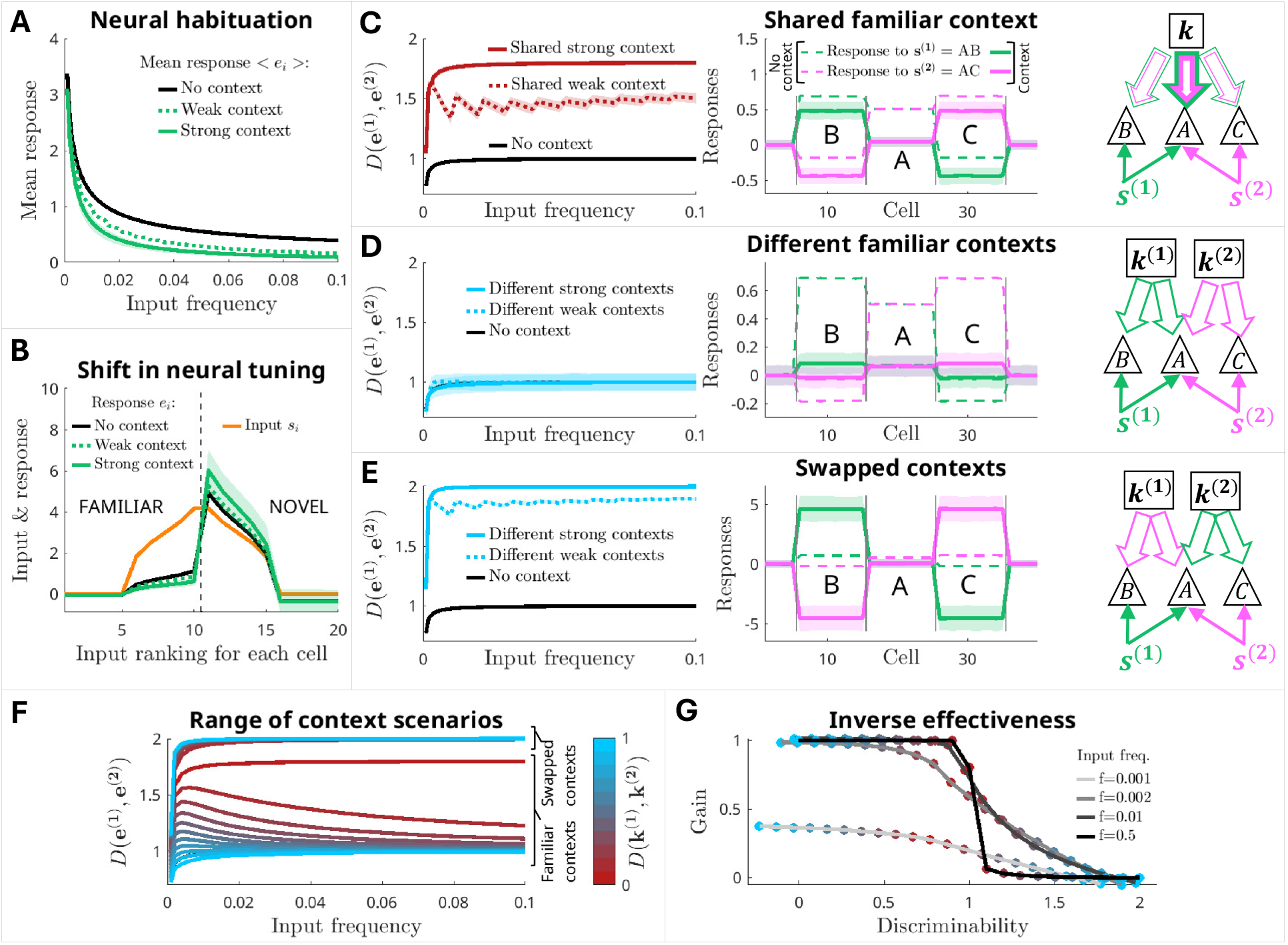
New theoretical predictions for multimodal interactions. **(A)** Habituation is more pronounced in the presence of cross-modal context (green) than in unimodal processing (black) and increases with input-context association strength (dotted line: weak; solid line: strong). **(B)** E cells sharpen their selectivity for novel stimuli in the presence of cross-modal contexts (green) relative to unimodal processing (black); see Methods for details. **(C–E)** The magnitude of pattern separation depends on the specific associations between stimuli and contexts. Dotted/solid lines: weak/strong stimulus-context association, as in (A–B). **(C)** Two stimuli (***s***^**(1)**^ = (*A, B*) and ***s***^**(2)**^ = (*A, C*)) are associated with the same context ***k***. Left: compared to the case without context (black), the cosine distance between responses is significantly higher. Middle: E-cell responses to input components *A, B*, and *C*, with context (solid lines) and without context (dashed lines). The shared context more strongly predicts (and suppresses) responses to the common component *A*. Mean and IQR across different stimulus examples are shown. Right: Schematic of the effects induced by feedback on E responses (triangles). Same-color arrows indicate concurrent activation; line width indicates inhibitory feedback strength. Compare with Middle. **(D–E)** Same as in (C), but here the stimuli are associated with orthogonal contexts, ***k***^**(1)**^ and ***k***^**(2)**^, respectively. **(D)** When responses to ***s***^**(1)**^ and ***s***^**(2)**^ are assessed in their familiar contexts, the cosine distance remains unchanged compared to the case without context, since the contexts predict (and uniformly suppress) responses to both overlapping (*A*) and non-overlapping (*B* or *C*) components. **(E)** When responses are assessed in the alternative (unfamiliar) contexts, the cosine distance is maximized, since each context continues to predict (and suppress) the input components with which it was previously associated, enhancing contrast between representations. **(F)** Pattern separation of E-cell responses when stimuli are associated with contexts exhibiting intermediate cosine distances (colorbar). **(G)** Inverse effectiveness: the discriminability gain from contextual feedback (y-axis) decreases as the discriminability of independent representations (x-axis) increases. The sensory stimuli are held constant while the distance between contexts is progressively varied (colors as in (F)). Results for different input frequencies are shown (legend). At high frequencies, the network exhibits a sharp transition between maximal gain and no gain at an intermediate level of discriminability.

#### 2. Inverse-effectiveness

Another consequence of the theory is a form of “inverse effectiveness”, where cross-modal feedback enhances the discriminability of sensory neural representations more strongly when the discriminability of representations within each modality (without feedback interaction ***F*** ) is weaker and sensory information is most ambiguous. This is illustrated in three example scenarios (Fig. 5C–E).

First scenario, corresponding to intermediate sensory ambiguity (Fig. 5C): two similar unimodal inputs (e.g., two odor stimuli, ***s***^**(1)**^ = (*A, B*) and ***s***^**(2)**^ = (*A, C*) with overlapping component *A*) are always encountered in the same context, eliciting similar responses ***k*** from other senses (e.g., vision [15]). The complete overlap between the contexts and the similarity of ***s***^**(1)**^ and ***s***^**(2)**^ create some level of sensory ambiguity in the representations, in the absence of feedback interactions between modalities. When the feedback interactions (***F*** ) are activated, the shared context entirely predicts the overlapping input component (*A*) because this component is always present in that context. On the contrary, the components that are specific to each input (*B* and *C*) are not always present in that context and are less strongly predicted. As a consequence, the cross-modal feedback predominantly suppresses responses to *A*. This effect sharpens the selectivity of neural representations for each input and increases pattern separation.

Second scenario, corresponding to low sensory ambiguity (Fig. 5D): the two unimodal inputs (***s***^**(1)**^ and ***s***^**(2)**^) are always encountered in two different contexts (***k***^**(1)**^ and ***k***^**(2)**^, respectively). Here, the lack of overlap between the contexts reduces the level of sensory ambiguity compared to the first scenario. In this case, when the feedback interactions (***F*** ) are activated, each context equally predicts (and uniformly suppresses) both overlapping and non-overlapping input components, leaving representation selectivity and pattern separation unchanged compared to the case with no feedback.

Third scenario, corresponding to high sensory ambiguity (Fig. 5E): the unimodal inputs (***s***^**(1)**^ and ***s***^**(2)**^) are repeatedly encountered in distinct contexts (***k***^**(1)**^ and ***k***^**(2)**^, respectively); subsequently, the contexts are swapped, so that ***s***^**(1)**^ is suddenly found in ***k***^**(2)**^ and ***s***^**(2)**^ in ***k***^**(1)**^, creating a high level of sensory ambiguity. In this case, when the feedback interactions are activated, ***k***^**(1)**^ and ***k***^**(2)**^ predict ***s***^**(1)**^ and ***s***^**(2)**^, respectively. When the contexts are suddenly swapped, feedback inhibits the E cells tuned to the unexpectedly absent stimulus, maximizing contrast between the representations of ***s***^**(1)**^ and ***s***^**(2)**^.

When the contexts have intermediate correlations, i.e., they are neither identical nor orthogonal, the distance between sensory representations transitions smoothly between the three scenarios (Fig. 5F). In particular, the cosine distance between E-cell responses decreases with context distance when the stimuli are presented in their familiar contexts (interpolating between the first and the second scenarios), while it rapidly increases when the stimuli are presented in swapped contexts (interpolating between the first and the third scenarios).

In summary, the theory predicts that cross-modal feedback adapts early sensory representations to the demands of the overall multisensory context, while suppressing familiar, redundant, and non-specific information. Typically, feedback sharpens neural selectivity for novel stimuli and enhances discriminability between similar sensory representations.

The gain in pattern separation due to cross-modal feedback is inversely related to the discriminability of representations within each modality, meaning that larger gains occur when the representations are less discriminable (Fig. 5G, inverse effectiveness). All the predictions of Fig. 5 are falsifiable, suggest plausible functions for cross-modal feedback in sensory coding, and provide a coherent framework for guiding future experiments.

## III. DISCUSSION

We have developed an analytical theory of sensory coding for canonical circuits that integrate a local input stream from a given sensory modality with an extrinsic feedback stream from another sensory modality or from motor circuits. The theory naturally generalizes to the case of multiple feedback streams, which can be represented as distinct blocks within the correlation matrices ***C***_***K***_ and ***C***_***SK***_. In this class of circuits, we demonstrated that predictive codes emerge as a direct consequence of efficient coding, unifying these normative frameworks within a single coherent theory.

Below, we discuss how this framework differs from prior unifying approaches and existing formulations of predictive coding, how it relates to the broader experimental literature, how it generalizes under additional or more biologically realistic constraints, and we outline directions for future work.

### A. Distinction from other unifying normative frameworks

The equivalence we uncovered is between context-conditioned efficient coding and *retrospective* predictive coding, in which predictions are formed to explain away sensory inputs, leading to the encoding of information unexpected under the context. This equivalence does not extend to *prospective* predictive coding, which instead concerns encoding information predictive of future stimuli or task outcomes and therefore embodies a fundamentally different objective. Accordingly, our theory and conclusions should not be conflated with prior frameworks aimed at integrating sparse coding with *prospective* predictive coding, such as the Information Bottleneck framework and related formulations [81, 85]. These frameworks do not reveal equivalences between principles, but instead characterize trade-offs between compressing the past and predicting the future (i.e., prospective predictive coding), recovering distinct coding objectives in different parameter regimes.

### B. Algorithmic interpretation

The unified normative theory proposed here implies a neural algorithm, in which each circuit component serves a well-defined function: cross-modal feedback delivers optimal predictions of sensory inputs, the local E-I network implements the precision matrix, and excitatory neurons integrate these elements with feedforward sensory stimuli, encoding whitened mixtures of prediction errors. A further probabilistic interpretation arises when the energetic penalty constrains response fluctuations: in this regime, the squared amplitude of E-cell response fluctuations scales as the negative log-likelihood of prediction errors, connecting to a natural mathematical definition of “surprise”. These insights advance our understanding of the operations performed by multimodal canonical circuits, bridging computational and algorithmic interpretations of context-dependent sensory coding.

### C. Distinction from other interpretations of retrospective predictive coding

Previous work has interpreted retrospective predictive coding as redundancy reduction within unimodal sensory streams [52, 115], as approximate Bayesian inference in hierarchical generative models [71–75, 77, 78], or as learning-based predictive mechanisms, often in sensorimotor circuits, where experience and error signals shape mappings from movement- or context-related activity to sensory predictions [29, 30, 36, 79]. Those formulations of predictive coding differ from ours in several important aspects.

First, in [52, 115], predictions are derived from statistical regularities within a single sensory stream, and redundancy is reduced by subtracting these internally generated predictions through local interactions among sensory neurons. By contrast, our theory explicitly factorizes redundancy reduction into two complementary processes: a local decorrelation mechanism operating within modalities and a predictive mechanism integrating information across modalities. The former is achieved through a precision-based whitening operation implemented via lateral effective connectivity between E cells, while the latter arises from extrinsic feedback and captures associations across different sensory streams or between motor commands and their sensory consequences. This decomposition of redundancy reduction into intrinsic and extrinsic processes establishes a previously unrecognized equivalence between predictive coding and context-conditioned efficient coding in multimodal networks.

Second, in classical Bayesian predictive coding [71–75, 77, 78], feedback predictions are generated by latent variables within a generative model, whose states are inferred from sensory data through iterative updates driven by prediction errors. By contrast, in our framework, contextual signals are externally supplied conditioning variables rather than latent causes to be inferred. Accordingly, the states of context neurons are not updated by bottom-up prediction-error signals in an online inference loop.

This perspective naturally accommodates the strong asymmetries observed in many crossmodal and sensorimotor interactions. Although reciprocal connections are widespread between cortical areas [125], opposing pathways can differ substantially in strength and do not necessarily participate in a closed inference loop. For example, auditory-to-visual connections are generally stronger than the reciprocal visual-to-auditory ones [6, 9, 126]; when influences from vision to audition are present, they often belong to distinct functional circuits with different roles [116, 117]. Similarly, several lines of evidence suggest that vision can strongly modulate olfactory perception without comparably strong reciprocal effects [7, 11, 15]. Another example is provided by motor efference copies, or corollary discharges. These signals are typically generated by motor command circuits and transmitted to sensory and other areas through dedicated pathways branching from the motor system. Experimental studies show that these pathways convey predictive information about impending movements or their sensory consequences, influencing sensory processing [34, 95, 127]. While in some cases sensory signals can reciprocally influence motor programs, for example during motor learning [36], in many other cases they do not alter the motor commands that generate the movement, such as when walking predicts footstep sounds but these sounds do not, in turn, alter walking.

These observations highlight that directional predictive interactions can arise without requiring a reciprocal inference loop in which prediction errors iteratively update latent state variables, as assumed in classical Bayesian predictive coding [71–75, 77, 78].

Recent experimental work, including [29, 30, 36, 79], emphasizes learning-based predictive mechanisms in which prediction errors act as teaching signals that can sometimes update internal models, i.e., mappings from movement-related signals to sensory predictions, rather than latent state variables. Our theory is consistent with this mechanistic interpretation, as prediction errors may drive Hebbian learning of the connectivity ***F*** that maps ***k*** to ***ŝ***. At the same time, it advances this perspective in two key ways. First, while previous models were mechanistic or descriptive, our theory provides a normative derivation of these effects based on context-conditioned efficient coding. Second, it links this normative account to an explicit algorithmic implementation in which predictive signals give rise to a precision-structured prediction-error code. In this framework, prediction errors are not merely teaching signals, but emerge as the normative code for efficient information transmission in the presence of context.

### D. Multisensory and sensorimotor effects from context-driven input compression

The cross-modal effects studied here have largely lacked a unified normative explanation [3, 4, 14, 15, 29, 30, 36, 37, 79]. Our theory shows that these effects can emerge directly from context-driven input compression. This simple normative principle provides a unified explanation of *why* a broad range of multisensory and sensorimotor interactions arise, including audiovisual [14], visual–olfactory [15], auditory–somatosensory [3, 4], audiomotor [35, 37], somatosensory–motor [40, 41, 95], and vestibular–motor effects [42, 43, 118]. Moreover, unimodal sensory coding emerges as a special case of the same framework when contextual feedback is absent. In this limit, the theory accounts for classical unimodal phenomena such as neural habituation [99–108], pattern separation [102, 108–110], and range adaptation [96–98, 119–121].

### E. Relation to mechanistic models

This normative perspective complements bottom-up mechanistic models that investigate *how* neuroplasticity implements adaptive computations [128], including habituation to familiar stimuli and amplification of novelty responses [129–131], competitive interaction effects [132], and the adaptive shaping of receptive fields [133] via synaptic plasticity, enhanced or reduced pattern separation via structural plasticity [134, 135], and whitening via gain modulation [136] or via combinations of gain modulation and synaptic plasticity [69]. Additional support for our framework comes from mechanistic modeling in olfaction, which shows that the granule cell activation patterns and receptive fields predicted by our theory can emerge through Hebbian plasticity [15] and adult neurogenesis [137]. The detailed convergence between our top-down normative framework and bottom-up models of neuroplasticity will be explored further in future work.

### F. A new form of inverse effectiveness

Given the relative scarcity of experiments on multimodal interactions across the senses, generating new falsifiable predictions is crucial for guiding future research in this area. Our framework yields a range of testable predictions, including a new form of inverse effectiveness, where feedback most effectively enhances the separation (decodability) of upstream sensory representations when information from each modality is most ambiguous. This prediction is broadly consistent with other forms of inverse effectiveness observed in experiments, which show that interactions between the senses are stronger when the effectiveness of unimodal stimuli is weaker [17, 22, 138]. However, previous experiments have primarily focused on the amplitudes of sensory responses, whereas our prediction concerns the relative relationships (distances) between sensory representations.

### G. Generality of the core principles and immediate extensions

Our analysis relies on simplifying assumptions, such as linear responses, Gaussian input statistics, quadratic (*L*_2_) metabolic costs, and high signal-to-noise, which enable an exact, closed-form solution. While these assumptions are not strictly biologically realistic [55, 58, 59], they primarily provide analytical tractability rather than defining the core principle of the theory. We have shown that our framework captures a broad range of experimentally observed phenomena, supporting its biological relevance. Here, we provide conceptual arguments suggesting that the equivalence between context-conditioned efficient coding and predictive coding, and the corresponding high-level mapping between circuit architecture and computation, do not depend on these specific assumptions, but instead follow from more general principles. We also discuss which aspects of the implementation are expected to depend on the underlying biological constraints.

Previous work has shown that biologically plausible constraints on neurons and circuits can uniquely select efficient coding solutions that remain degenerate under linear-Gaussian-*L*_2_ assumptions [58, 59, 139]. We have demonstrated that our key results apply across this entire class of degenerate solutions and are therefore invariant to transformations such as rotations and reflections. More generally, a robust consequence of context-conditioned efficient coding is that a circuit should remove the component of the input that is predictable from contextual signals in a reliable manner, and efficiently represent the remaining conditional uncertainty. Under architectures with distinct extrinsic and local pathways, this principle should naturally give rise to a functional decomposition in which extrinsic pathways processing contextual information convey predictions that subtract context-predictable structure, while local recurrent circuitry reduce redundancy in the residual representation. This principle is expected to persist beyond the specific assumptions considered here, whereas the form of the optimal predictor and the nature of residual recoding depend on neural and circuit constraints, input statistics, and noise conditions.

In general, feedback can be understood as computing a conditional estimator, 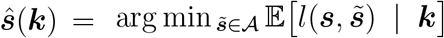, where the admissible class *A* is constrained by circuit and neural response properties, and the loss *l* specifies how prediction errors are penalized. The choice of metabolic cost function in the Efficiency objective and the input statistics determine which aspects of prediction errors are most costly to represent, i.e., the specific form of the loss *l* and, consequently, which estimator is optimal. In the linear-Gaussian-*L*_2_ case, this leads exactly to the conditional mean or the best linear projection of ***s*** onto ***k*** as the optimal predictors, when metabolic cost penalizes squared fluctuations or squared responses, respectively; under alternative regimes (e.g., heavy-tailed inputs or *L*_1_ costs favoring sparse activity), other estimators such as the conditional median or more general nonlinear functions of context are expected to emerge.

Under this broader view, the notion of “whitening” as optimal residual coding is exact under Gaussian assumptions, where second-order statistics fully characterize the distribution. For non-Gaussian residuals, efficient coding is expected to require reducing higher-order dependencies, leading to representations resembling sparse coding, independent component analysis, or nonlinear normalization. These generalizations are not derived here in full generality and remain an important direction for future work, but are expected to follow from previous results in statistical estimation and efficient coding [55, 58, 139, 140].

Finally, noise affecting both sensory inputs and contextual signals modifies but does not invalidate the decomposition into an extrinsic predictive component and a local residual coding component. In noisy settings, it is expected that the optimal predictor corresponds to a noise-aware conditional estimator, and the residual reflects both unpredictable signal fluctuations and noise variability conditional on context. Noise is therefore expected to shape, but not abolish, both prediction and residual coding, potentially shifting the balance between feedback and local processing [141], as well as between compression and redundancy, the latter becoming advantageous for reliable information transmission.

A full treatment of these extensions, particularly in nonlinear, non-Gaussian, and noisy regimes, is left for future work.

### H. Open problems and future directions

In conclusion, our work represents an initial step toward establishing a conceptual framework for understanding neural coding in multimodal networks. Sensory systems exhibit diverse circuit motifs, many of which go beyond the canonical networks studied here, and negative prediction-error neurons – which increase rather than decrease their activity when a sensory stimulus is smaller than predicted – have also been observed, for example in vision [26, 28, 29, 32, 33, 142]. Other multisensory and sensorimotor interactions exist that are not covered in this study. Additionally, other types of contextual signals, such as stimulus valence or behavioral variables, are also transmitted to the sensory periphery via feedback pathways [143]. Our theory provides a normative foundation that can be extended to address these scenarios. First, this framework can be generalized to study sensory coding in more structured networks by expanding and constraining the model in Fig. 1A. This includes incorporating different cell types and connectivity motifs, such as feedforward, feedback, lateral inhibition, and disinhibitory motifs [144]. Second, different coding hypotheses may be formulated by modifying the coding functional in Fig. 1D – for instance, to integrate task-oriented objectives, as proposed in [81, 120, 145–148], or to explore broader normative principles that balance general-purpose information transmission with task-specific behavioral goals. Advancing theory in these directions is crucial for understanding the roles of different circuit motifs, feedback pathways, and plasticity mechanisms in optimizing neural codes across sensorimotor and decision-making networks.

## IV. METHODS

### A. Multimodal canonical circuit model: reduction to effective description

We define the vectors ***e*** = (*e*_1_, …, *e*_*N*_ ) as the responses of excitatory neurons, ***i*** = (*i*_1_, …, *i*_*Q*_) as the responses of inhibitory neurons, ***s*** = (*s*_1_, …, *s*_*N*_ ) as the feedforward sensory inputs, ***k*** = (*k*_1_, …, *k*_*P*_ ) as the cross-modal feedback inputs, and ***J***^*XY*^ as the matrix of synaptic weights from every neuron in class *Y* ∈ (E, I, K) to every neuron in class *X* ∈ (E, I, K). We define the following linear response model (Fig. 1A):

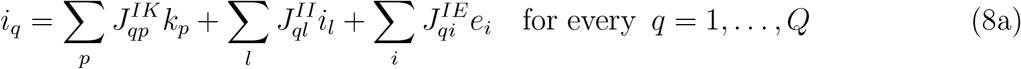

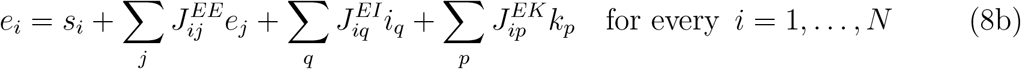

where all entries of ***J***^*EE*^ and ***J***^*IE*^ are positive, all entries of ***J***^*II*^ and ***J***^*EI*^ are negative, and the remaining weight matrices are unconstrained. The model can be rewritten in a compact form as:

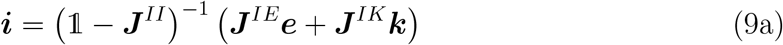

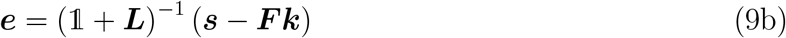

where ***L*** and ***F*** are the matrices of effective lateral connections between E neurons and effective feedback connections from K to E neurons, respectively, and 1 denotes the identity matrix. These effective connections are functions of the synaptic weights:

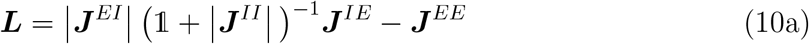

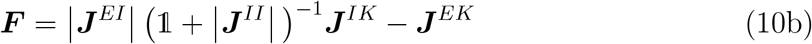

From Eq. (9b), we obtain the multimodal responses ***R*** = (***e, k***) as a function of the multimodal inputs ***I*** = (***s, k***):

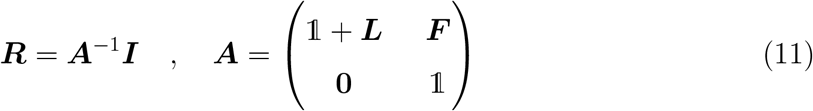

where ***R*** and ***I*** are column vectors to ensure dimensional consistency. We consider canonical circuits where effective lateral connections are symmetric, i.e., ***L*** = ***L***^*T*^ . This symmetry constraint allows us to derive a closed-form solution to the efficient coding problem.

### B. Derivation of the optimal connectivity for multimodal efficient coding

#### 1. Multimodal Efficiency functional

For general noise conditions, we can define multimodal Efficiency as

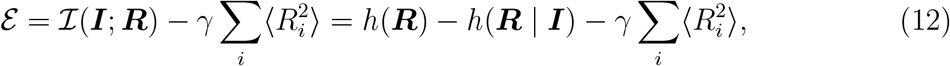

where *h*(·) denotes differential entropy (since ***R*** is continuous).

Here, we consider a regime of vanishing noise but finite readout precision. In the zero-noise (deterministic) limit, the mutual information between continuous variables diverges or becomes ill-defined, unless a finite-precision readout is imposed. Following standard approaches [149], we therefore consider a quantized response ***R***_Δ_ = *Q*_Δ_(***R***), where *Q*_Δ_ : ℝ^*d*^ → ℤ^*d*^ (*d* = *N* + *P* ) is a quantization map associated with a partition 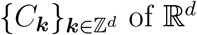, such that *Q*_Δ_(***r***) = ***k*** iff ***r*** ∈ *C*_***k***_. We assume the bins {*C*_***k***_} to be hypercubic with side length Δ and therefore equal volume, vol(*C*_***k***_) = Δ^*d*^ (for simplicity; more generally, all bins share the same volume up to a constant factor). In the deterministic limit ***R*** = *f* (***I***), we have H(***R***_Δ_ | ***I***) = 0 and:

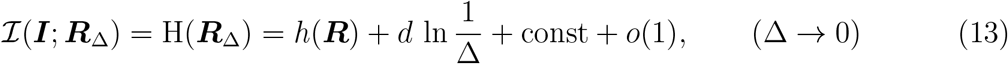

where H(·) denotes discrete entropy. The term *d* ln(1*/*Δ) depends only on resolution, while const depends on bin geometry and log base but not on the encoding; therefore, for fixed Δ, maximizing *I* (***I***; ***R***_Δ_) is equivalent to maximizing *h*(***R***) [149].

In conclusion, we can drop the constant terms (which do not affect the optimization) and define multimodal Efficiency as:

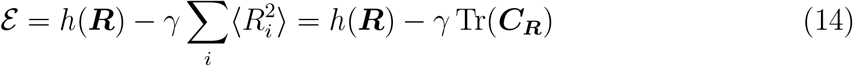

The energetic cost term is given by the trace of the second-moment matrix of the responses, 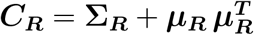.

We approximate the multimodal input distribution as a multivariate Gaussian, *p*(***I***) = N(***µ*, Σ**). This represents the maximum-entropy (i.e., least-constrained or least-biased) distribution that preserves all input means and pairwise correlations, allowing us to understand how canonical networks should adapt to these low-order statistics to efficiently encode stimuli. Under this approximation, the responses are also normally distributed, *p*(***R***) = N(***µ***_***R***_, **Σ**_***R***_ ), with mean ***µ***_***R***_ = ***A***^−1^***µ*** and covariance **Σ**_***R***_ = ***A***^−1^**Σ** (***A***^−1^)^*T*^ . We can thus analytically express *E* as a function of the circuit effective connectivity, summarized by ***A***, the input covariance **Σ**, and the second-moment matrix of the inputs (referred to simply as the input correlation matrix), ***C*** = **Σ** + ***µµ***^***T***^ :

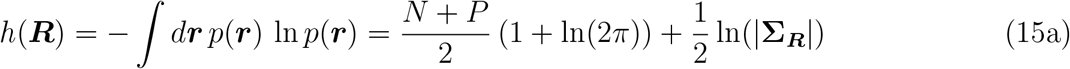

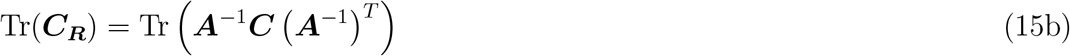

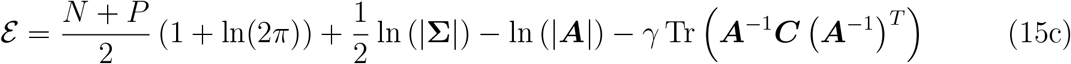

#### 2. Stationary point

We compute the derivatives of E with respect to the effective connections *L*_*ij*_ and *F*_*ip*_. These are defined as:

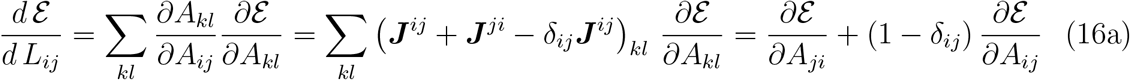

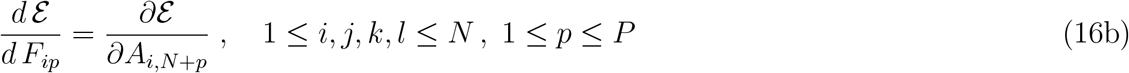

where ***J***^*ij*^ is the matrix with element (*i, j*) equal to 1 and zeros elsewhere, so that (***J***^*ij*^)_*kl*_ = *δ*_*ik*_ *δ*_*jl*_ . The sum over partial derivatives in Eq. (16a) arises due to the symmetry of ***L***. Using matrix algebra, we compute the partial derivatives 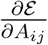, for 1 ≤ *i* ≤ *N* and 1 ≤ *j* ≤ *N* + *P* :

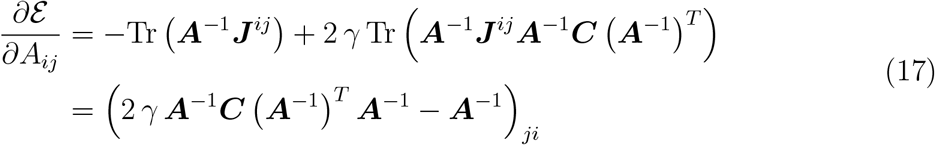

We use the Schur complement of the bottom-right block of ***A*** to invert ***A***, and the symmetry of ***L*** to transpose ***A***^−1^. We obtain:

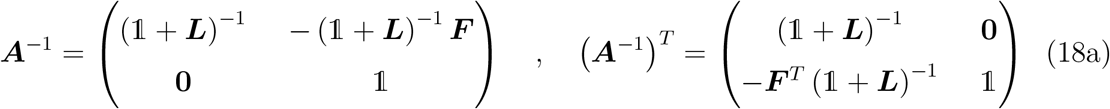

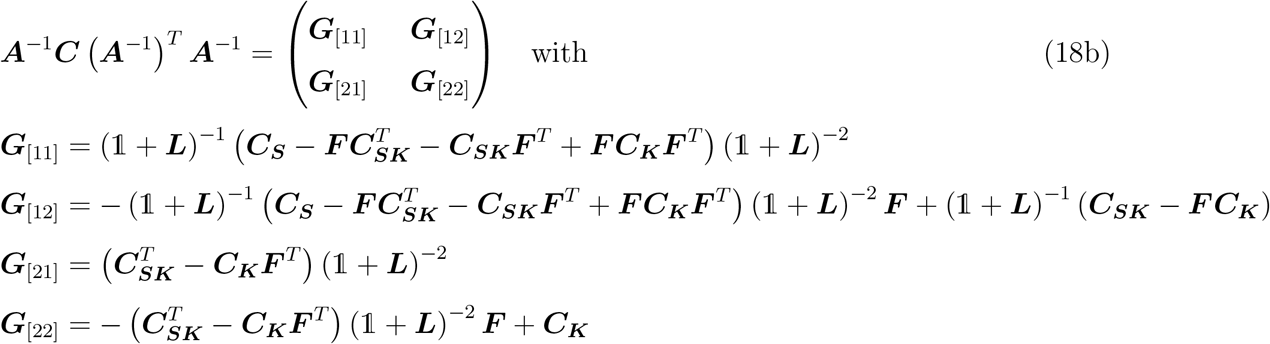

Here and in what follows, the square-bracket subscripts denote matrix blocks. Substituting Eqs. (18) into (17) and Eqs. (17) into (16), we obtain:

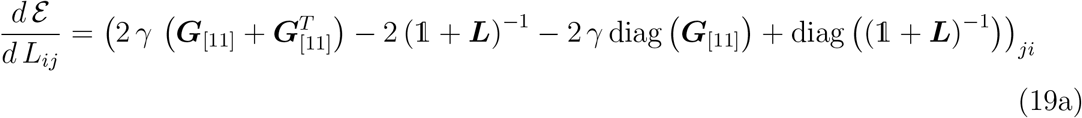

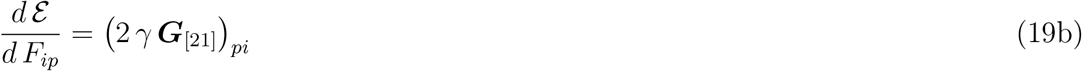

Solving 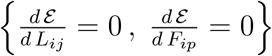 for all 1 ≤ *i, j* ≤ *N* and 1 ≤ *p* ≤ *P*, we obtain the following stationary point:

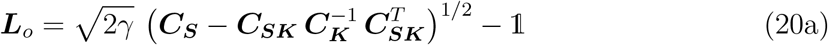

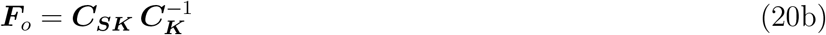

Below, we prove three facts: (1) this stationary point maximizes multimodal Efficiency; thus, it is an optimal solution; (2) it is one particular solution within a class of degenerate solutions, all of which have the same Efficiency; (3) within this class, the solution given by Eqs. (20) is the one for which the responses ***R*** are most similar to the inputs ***I*** (i.e., the most local encoding).

#### 3. Maximization of multimodal Efficiency

To prove that the stationary point 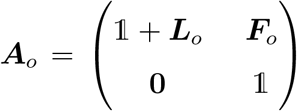 maximizes multimodal Efficiency, we compute the Hessian of *E* at ***A***_*o*_ and demonstrate that it is negative definite (i.e., all its eigenvalues are negative).

The four blocks of the Hessian, corresponding to the second-order partial derivatives of E with respect to different combinations of ***L*** and ***F*** elements, are defined as follows. The differences between the blocks arise from the fact that ***L*** is symmetric, whereas ***F*** is not:

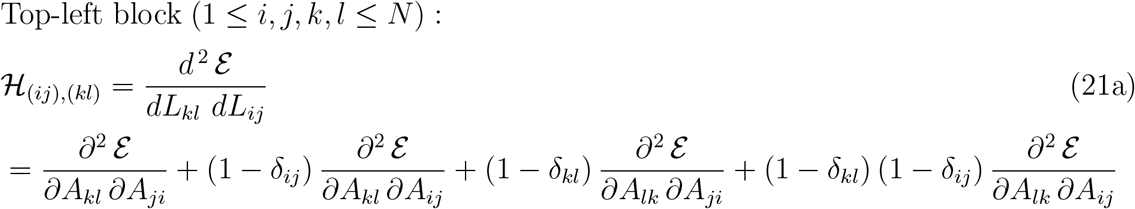

Top-right block (1 ≤ *i, j, k* ≤ *N, N < l* ≤ *N* + *P* ) :

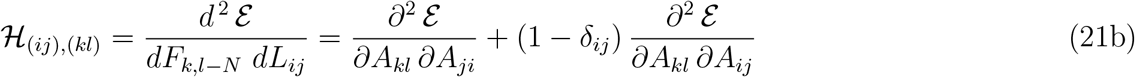

Bottom-left block (1 ≤ *k, l, i* ≤ *N, N < j* ≤ *N* + *P* ) :

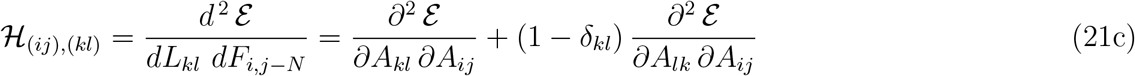

Bottom-right block (1 ≤ *i, k* ≤ *N, N < j, l* ≤ *N* + *P* ) :

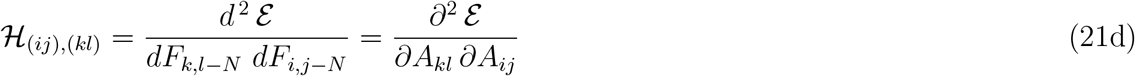

We only need to compute 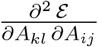, as the other second-order partial derivatives are obtained by simply permuting the indices:

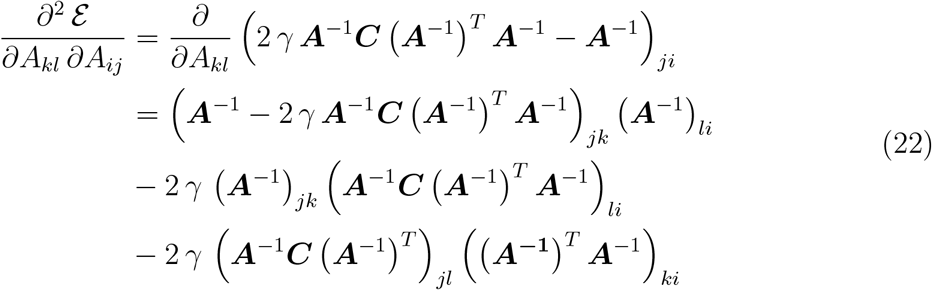

At the stationary point, the first term in Eq. (22) vanishes and the matrix products simplify as follows:

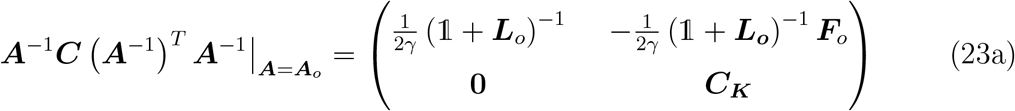

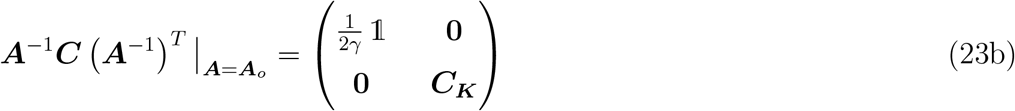

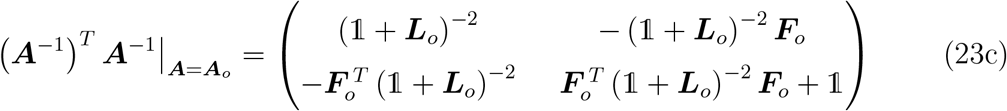

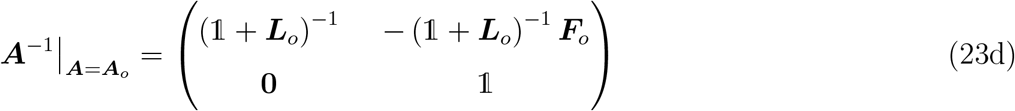

Depending on the range of the indices *i, j, k, l*, we need to consider different blocks within the matrices in Eqs. (23). Substituting the corresponding matrix blocks from Eqs. (23) into Eq. (22), we obtain:

Top-left block (1 ≤ *i, j, k, l* ≤ *N* ) :

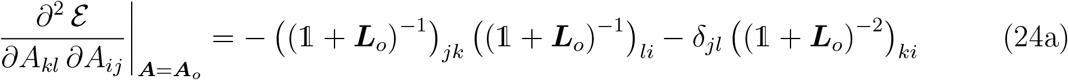

Top-right block (1 ≤ *i, j, k* ≤ *N, N < l* ≤ *N* + *P* ) :

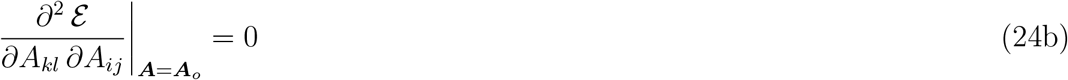

Bottom-left block (1 ≤ *k, l, i* ≤ *N, N < j* ≤ *N* + *P* ) :

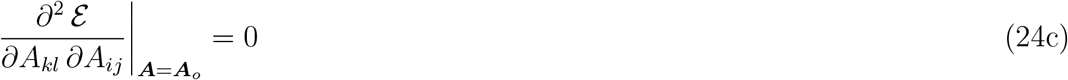

Bottom-right block (1 ≤ *i, k* ≤ *N, N < j, l* ≤ *N* + *P* ) :

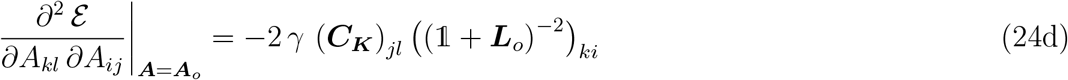

Finally, we sum the second-order partial derivatives as defined in Eqs. (21), using Eqs. (24) for the corresponding blocks. We obtain:

Top-left block (1 ≤ *i, j, k, l* ≤ *N* ) :

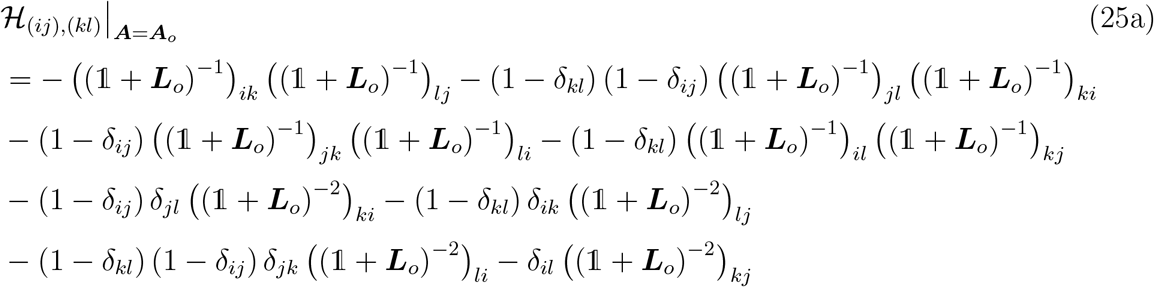

Off-diagonal blocks (1 ≤ *i, j, k* ≤ *N, N < l* ≤ *N* + *P* or 1 ≤ *k, l, i* ≤ *N, N < j* ≤ *N* + *P* ) :

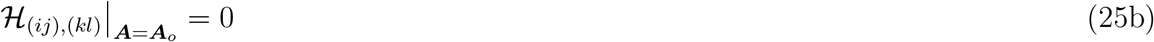

Bottom-right block (1 ≤ *i, k* ≤ *N, N < j, l* ≤ *N* + *P* ) :

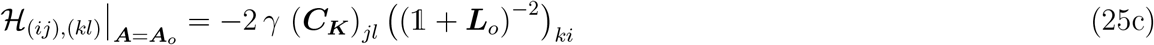

Eqs. (25) provide the full Hessian matrix at the stationary point. Since it is a block diagonal matrix, its eigenvalues are the eigenvalues of the two diagonal blocks. Thus, we need to prove that the top-left and bottom-right blocks are negative definite. These blocks can be rewritten as follows:

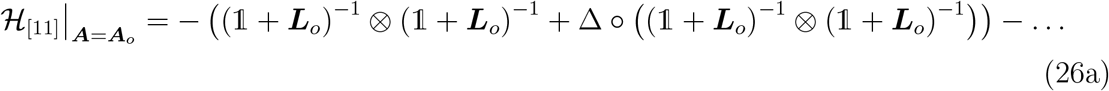

… individually asymmetric terms

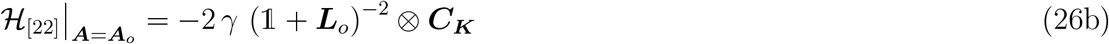

where Δ_(*ij*),(*kl*)_ = (1 − *δ*_*ij*_) (1 − *δ*_*kl*_), and ⊗, ◦ denote the Kronecker and Hadamard products, respectively.

The block 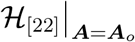 is negative definite because: (a) ***C***_***K***_ is an invertible correlation matrix and therefore positive definite; (b) 2 *γ* (1 + ***L***_*o*_)^−2^ = ***P*** = (***C***^−1^)_[11]_ is the top-left block of the precision matrix (or inverse correlation matrix); for any invertible correlation matrix, ***C***^−1^ is positive definite, and by Sylvester’s criterion, any principal submatrix of a positive definite matrix is also positive definite; thus, ***P*** is positive definite; (c) the Kronecker product of two positive definite matrices is positive definite; (d) the negative sign in front ensures that 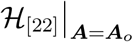 is negative definite.

To analyse the block 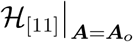, we use a hybrid analytical-numerical approach. The first two terms in Eq. (26a) correspond to the first two terms in Eq. (25a), which are symmetric, i.e., invariant under the permutations of indices *i* ⇔ *k, j* ⇔ *l*. These terms can be analyzed analytically: (a) 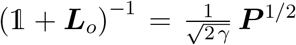 is a power of a positive definite matrix and is therefore positive definite; (b) the Kronecker products of such matrices are also positive definite; (c) Δ is positive semidefinite because ***x***^*T*^ Δ***x*** = ∑_*ij,kl*_ *x*_*ij*_ (1 − *δ*_*ij*_) (1 − *δ*_*kl*_) *x*_*kl*_ = (∑_i≠ *j*_^*x*^_*ij*_)^2^ ≥ 0 for every non-zero vector ***x***; (d) the Hadamard product of a positive semidefinite matrix and a positive definite matrix is positive semidefinite; (e) the sum of a positive definite matrix (first term) and a positive semidefinite matrix (second term) is always positive definite; (f) thus, the negative sum of these two terms is negative definite. The additional terms in Eq. (25a) are not individually symmetric (although their sum is symmetric, as expected). Thus, they are not analytically tractable. To complete the proof of the negative definiteness of 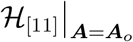 (and the overall Hessian matrix), we numerically computed the eigenvalues of 20, 000 Hessian matrices randomly generated using Eqs. (25) with ***L***_*o*_ given by Eq. (20a). The blocks of the correlation matrix ***C***, i.e., ***C***_***S***_, ***C***_***SK***_, and ***C***_***K***_, were obtained for inputs randomly sampled from Gaussian distributions. We used *N* = 50 units to represent the feedforward inputs (***s***), *P* = 50 units to represent the feedback inputs (***k***), and *T* = 1000 observations for each input. In the first batch of 10, 000 numerical simulations, the inputs were drawn from different Gaussian distributions: *I*_*i*_ ∼ N(*µ*_*i*_, *σ*), where *i* denotes the unit. The means varied across units and were themselves sampled from a Gaussian distribution: *µ*_*i*_ ∼ N(*µ*_0_, *σ*_0_). Both *σ*_0_ and *σ* were varied across simulations to generate different random matrices, allowing for variations in both the amplitude and heterogeneity of pairwise correlations. In all 10, 000 simulations, all eigenvalues of the resulting Hessian matrices were negative up to machine precision. In the second batch of 10, 000 simulations, the inputs were constructed similarly to the analyses in Figs. 4, 5: 20% of input units of each type were active at a rate of 0.02 (i.e., in 20 out of *T* = 1000 observations). Inputs were drawn from a Gaussian distribution: *I*_*i*_ ∼ N(*µ, σ*), with mean *µ* = 3 and standard deviation *σ* = 1. White Gaussian noise *η*_*i*_ ∼ N(0, *σ*_*n*_) with *σ*_*n*_ = 0.2 was added to all observations. Again, in all 10, 000 simulations, all eigenvalues of the resulting Hessian matrices were negative up to machine precision. Thus, we conclude that the stationary point given by Eqs. (20) maximizes multimodal Efficiency.

#### 4. Degenerate solutions

The determinant of **Σ**_***R***_ in Eq. (15a) and the trace of ***C***_***R***_ in Eq. (15b) are invariant under the transformation ***A***^−1^ → ***UA***^−1^, where ***U*** is an orthogonal matrix. Therefore, any encoding matrix of the form ***A***^′^ = ***AU***^*T*^ has the same Efficiency E. The matrices ***U*** and ***A***^′^ must satisfy the following equations, which ensure that ***U*** is orthogonal and that ***A***^′^ is an encoding matrix, i.e., it is consistent with the circuit structure, Eq. (11):

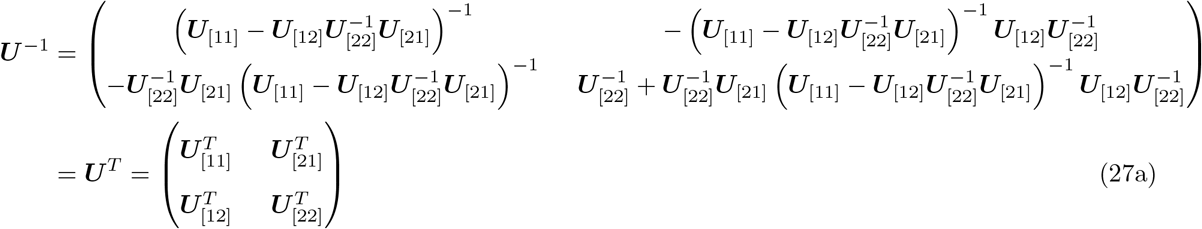

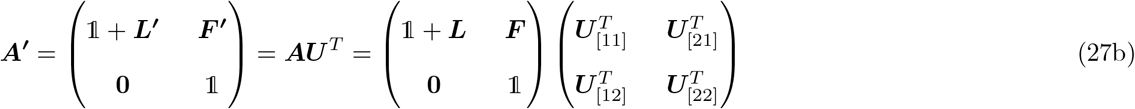

where we have used the Schur complement formula for inverting ***U*** . Renaming ***U***_[11]_ = ***Q***for simplicity and solving Eqs. (27a), we obtain:

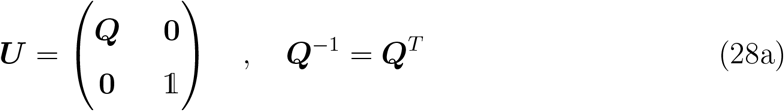

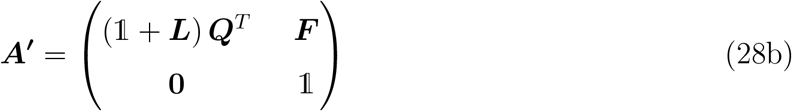

In conclusion, any lateral and feedback connectivity of the form

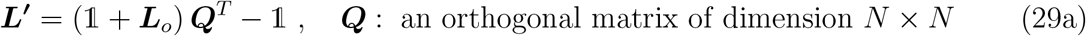

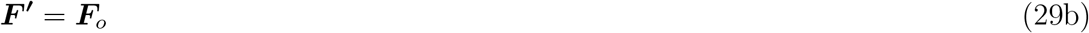

maximizes multimodal Efficiency. Eqs. (29) thus define a class of degenerate solutions. These solutions give rise to a class of representations ***e***^′^ that differ only by rotations or reflections:

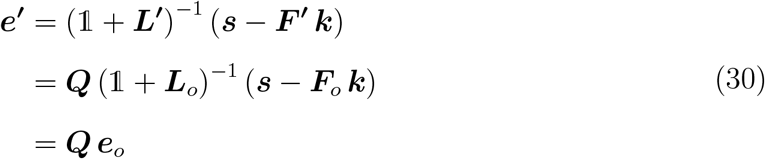

Here, ***e***_*o*_ = (1 + ***L***_*o*_)^−1^ (***s*** − ***F***_*o*_ ***k***) denotes the local responses, Eq. (9b), at the stationary point defined by Eqs. (20). Since ***Q*** is an orthogonal matrix, ***e***^′^ corresponds to a rotation or reflection of ***e***_*o*_. Thus, the geometry of the representation space remains unchanged across the entire class of degenerate solutions.

#### 5. Most local encoding

Within this class of degenerate solutions, we identify the most local encoding, i.e., the particular solution that minimizes the mean squared distance between inputs and responses. This corresponds to minimizing the mean squared distance between ***s*** and ***e*** (since ***k*** is a constant component in both the input and response vectors), subject to the constraint that ***Q*** is orthogonal. We thus seek the matrix ***Q*** that minimizes the following Lagrangian:

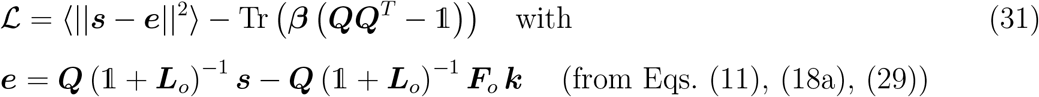

where ***β*** is the symmetric Lagrange multiplier matrix. The equations 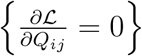 for all *i* = 1, …, *N* and *j* = 1, …, *N*, combined with Eqs. (20), lead to the following condition:

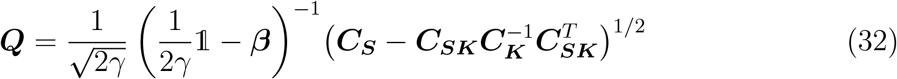

Using the orthogonality condition ***QQ***^*T*^ = 1, we obtain the Lagrange multiplier matrix:

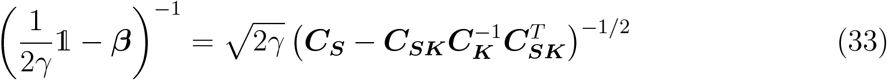

Combining Eqs. (33), (32), and (29), we obtain:

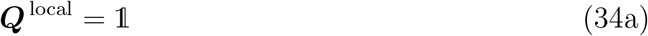

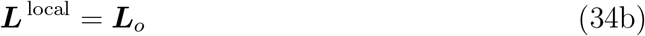

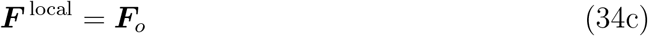

In conclusion, ***L***_*o*_ and ***F***_*o*_, as given in Eqs. (20), represent the unique connectivity solution that maximizes multimodal Efficiency and provides the most local encoding.

#### 6. Equivalence of joint and conditional multimodal Efficiency maximization

For arbitrary noise conditions, multimodal Efficiency is equivalent to:

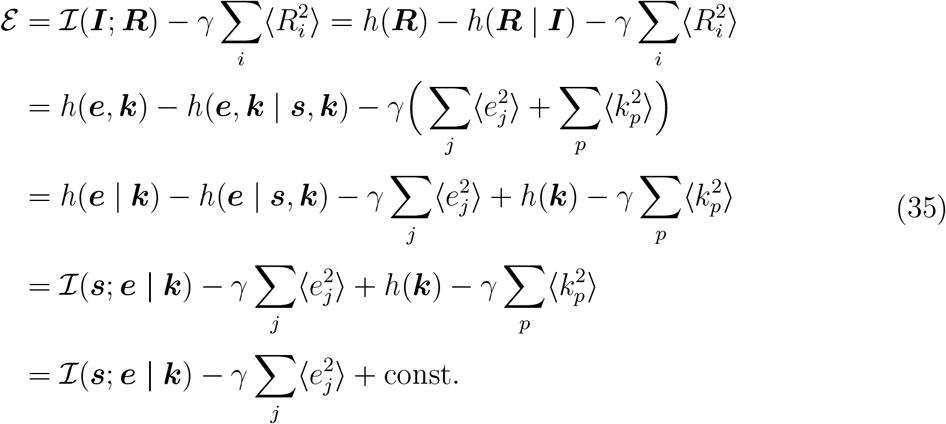

where 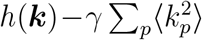 is constant with respect to the optimization parameters (***L*** and ***F*** ). Thus, for arbitrary noise, maximizing the mutual information between the multimodal inputs, ***I*** = (***s, k***), and the joint multimodal responses, ***R*** = (***e, k***), is equivalent to maximizing the mutual information between the local inputs, ***s***, and the local responses conditioned on the context, ***e*** | ***k***. Here, we consider the zero-noise limit, such that:

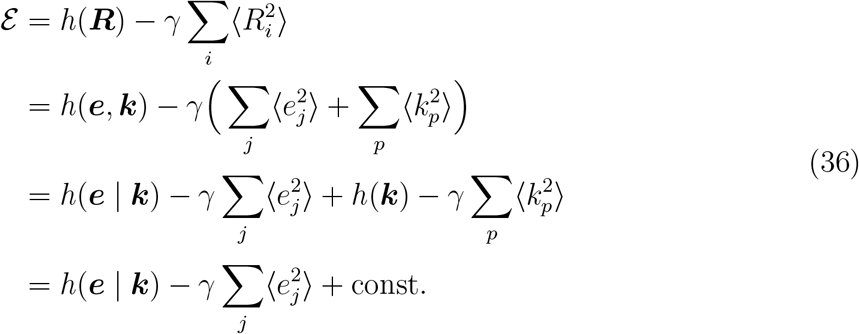

In conclusion, the problems of maximizing the entropy of joint multimodal responses and of maximizing the entropy of local responses conditioned on the context are equivalent. Under the same constraint on response amplitudes, both formulations give rise to the same class of degenerate solutions.

### C. Equivalence between multimodal (or context-conditioned) efficient coding and predictive coding

#### 1. Introductory roadmap

Here, we summarize the central equivalences between efficient and predictive coding; detailed mathematical derivations and proofs are provided in the subsequent sections of the Methods.

The central equivalences are:

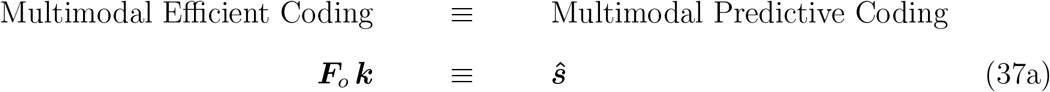

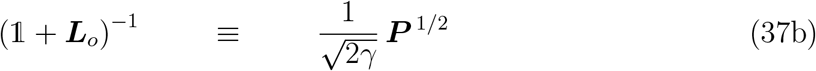

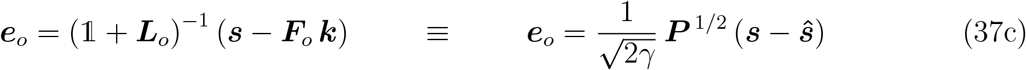

where ***L***_*o*_ and ***F***_*o*_ represent the optimal connectivity given by Eqs. (20). The left-hand side describes the most efficient encoding of local inputs under extrinsic context. The right-hand side describes the most predictive encoding: feedback transmits optimal linear predictions (***ŝ***), lateral connectivity encodes the precision matrix (***P*** ), and the excitatory neurons transmit mixtures of prediction errors (***e***_*o*_) downstream. These equivalences are derived in the next section.

Next, we derive the general properties of the predictive coding formula (right-hand side of Eq. (37c)).

Finally, we derive three element-wise interpretations of this formula that apply in special cases. The first two interpretations are equivalent and apply when the precision matrix exhibits a dominant-mode structure in which the eigenvalues of ***P*** cluster into two groups: *λ*_*k*_ ≈ *λ*_1_ (the largest eigenvalue) or *λ*_*k*_ ≪ *λ*_1_. In this regime, 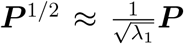, and each entry of the predictive coding formula can be expressed in the following equivalent forms:

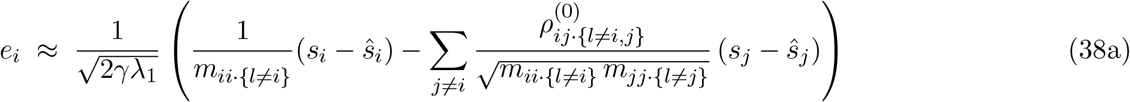

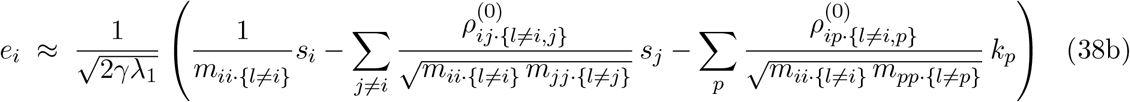

where *m*_*ii*·{*l*≠*i}*_ is the partial second moment of the *i*-th input and 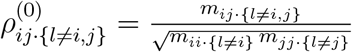 is the scale-normalized partial cross-moment between the *i*-th and *j*-th inputs. These represent, respectively, the residual uncentered second moment and a scale-normalized measure of residual cross-dependence after accounting for the linear effects of all other *l*-th inputs in (***s, k***).

The first interpretation, formula (38a), emphasizes that the circuit transmits precision-weighted, redundancy-reduced prediction errors. In this view, contextual feedback is used to compute optimal linear predictions (assuming zero intercept), ***ŝ***, of the local inputs ***s***, while lateral connectivity subtracts prediction-error components in proportion to their redundancy. The second – equivalent – interpretation, formula (38b), highlights that feedback inputs {*k*_*p*_} enter the same subtractive pathways as the feedforward inputs {*s*_*j*_}, *j*≠ *i*: through the circuit connectivity, both act to suppress redundant components in the transmission of each local input *s*_*i*_.

The third element-wise interpretation applies when the precision matrix is diagonally dominant, that is, ***P*** = ***D*** + ***E***, where ***D*** is diagonal, ***E*** is purely off-diagonal, and ***E*** is small relative to ***D*** in spectral norm. In this regime, the right-hand side of Eq. (37c) can be expressed in the following form:

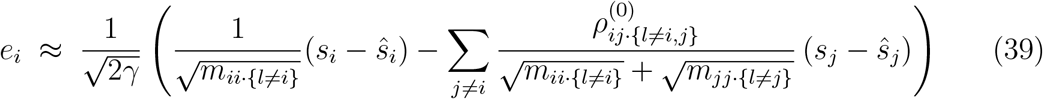

#### 2. Proof of equivalences (37)

The optimal predictions *ŝ*_*i*_ of *s*_*i*_ that can be linearly computed (with zero intercept) from 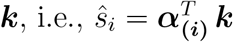, are obtained by finding the coefficient vectors ***α***_**(*i*)**_ that minimize the mean squared errors:

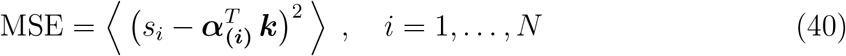

Solving 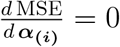 for every *i* = 1, …, *N*, we obtain the matrix of coefficients:

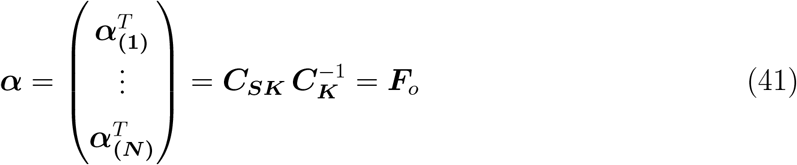

Thus, ***ŝ*** = ***α k*** = ***F***_*o*_ ***k***, which proves Eq. (37a).

Eq. (37b) is a direct consequence of Schur’s inversion formula for block matrices:

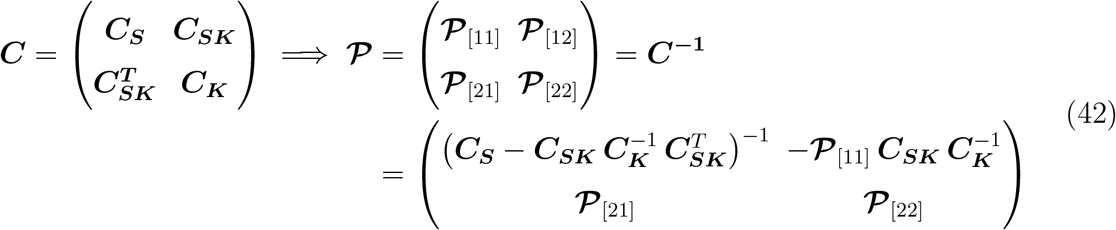

***P*** is the full precision matrix. Its top-left block, ***P***_[11]_ = ***P***, is the matrix that appears in the predictive coding formula, which we refer to simply as the precision matrix. Combining Eq. (20a) with Eq. (42), we obtain:

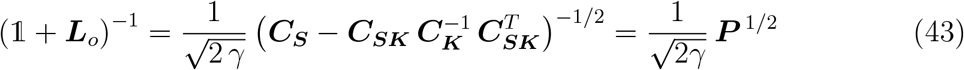

which completes the proof of Eq. (37b).

Eq. (37c) follows directly from Eqs. (37a) and (37b).

#### 3. General properties of the predictive coding formula, Eq. (37c)

Eq. (37c) implies that E neurons encode and transmit specific linear mixtures of prediction errors. This encoding has two key properties.

First, the mixtures conveyed by different neurons are mutually uncorrelated and have equal second moments, that is, 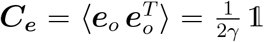. In other words, the representation is whitened with respect to the raw second-moment matrix ***C***_***ε***_ of the prediction errors ***ε*** = ***s*** − ***ŝ***.

Proof:

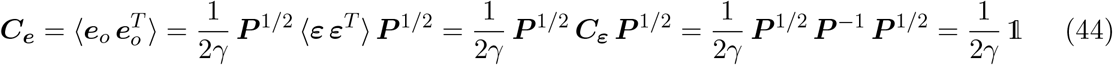

where we used:

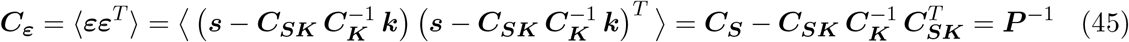

This property generalizes to the entire class of degenerate solutions ***e***^′^ = ***Q e***_*o*_, where ***Q*** is orthogonal.

Second, the Euclidean norm of the population response, ∥***e***_*o*_∥, represents the magnitude of the prediction-error vector ***ε*** = ***s*** − ***ŝ*** relative to its typical scale, namely 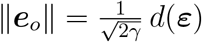, where 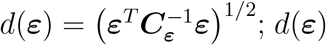 defines a Mahalanobis-like distance induced by the error second-moment matrix rather than the covariance matrix.

Proof:

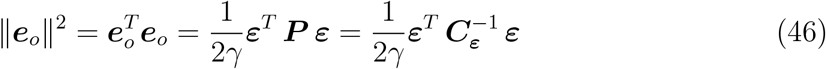

This property also generalizes to the entire class of degenerate solutions ***e***^′^ = ***Q e***_*o*_.

For a clean probabilistic interpretation see “Appendix: Generalization to a variance-based cost function”.

#### 4. Proof of formula (38a)

For precision matrices defined as ***P*** = **Σ**^−1^, a standard result in matrix analysis relates ***P*** to partial variances and partial correlations:

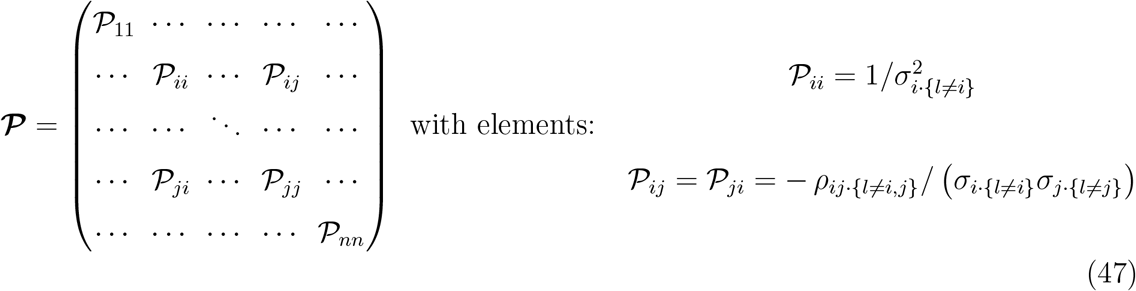

In this study, we use a slightly different definition, where ***P*** is the inverse of the raw second-moment matrix ***C*** = ⟨***II***^*T*^ ⟩, rather than the inverse of the covariance matrix **Σ** = ⟨***II***^*T*^ ⟩ − ⟨***I***⟩⟨***I***⟩^*T*^ . In the following sections, we prove that Eqs. (47) generalize to this setting after replacing the partial variances 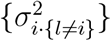 with partial second moments {*m*_*ii*·{l≠=*i*}_}, and the partial correlations {*ρ*_*ij*·{l≠=*i,j*}_} with scale-normalized partial cross-moments 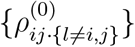.

Noting that *P*_*ij*_ = P_*ij*_ for 1 ≤ *i, j* ≤ *N*, we substitute these generalized expressions into the right-hand side of Eq. (37c), under the approximation 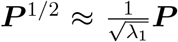. This leads to the formula (38a).

Equivalently, 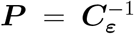, which yields an analogous interpretation in terms of partial second moments and partial cross-moments of the prediction errors.

##### Diagonal elements of the precision matrix

We consider the linear regression *with zero intercept* of *s*_*i*_ on ***z***_**(*i*)**_ = (***s, k***) \ *s*_*i*_ (i.e., the vector of all local and contextual inputs excluding *s*_*i*_). The residuals of this regression are 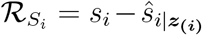, where 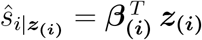 is the minimum mean squared error linear estimator of *s*_*i*_ given ***z***_**(*i*)**_ and zero intercept. We then define the partial second moment *m*_*ii*·{l≠=*i*}_ as the second raw moment of these residuals:

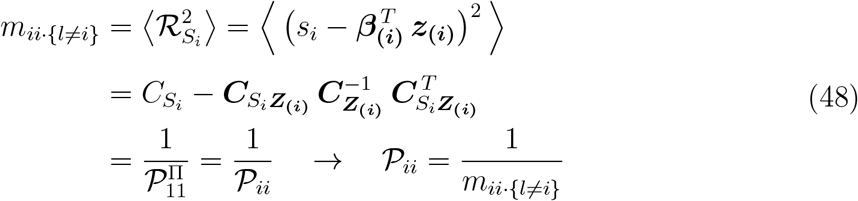

Here, we denote by 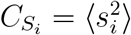 the scalar second moment of *s*_*i*_, by 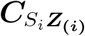 the row vector of second cross-moments between *s*_*i*_ and each element of ***z***_**(*i*)**_, and by 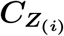 the matrix of second moments between the elements of ***z***_**(*i*)**_.

On the second line of Eq. (48), we use the fact that 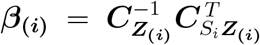, analogous to Eq. (41). On the third line, we first apply a permutation to the matrix ***C*** that swaps its *i*-th and first rows and columns: ***C***^**Π**^ = **Π *C* Π**, where **Π** is the symmetric permutation matrix that swaps these coordinates (defined as Π_*kl*_ = *δ*_*kl*_ − *δ*_*ki*_ *δ*_*li*_ − *δ*_*k*1_ *δ*_*l*1_ + *δ*_*ki*_ *δ*_*l*1_ + *δ*_*k*1_ *δ*_*li*_). Next, we partition ***C***^**Π**^ so that its top-left block consists only of its (1, 1) element, which corresponds to the (*i, i*) element of ***C***. We then apply the Schur complement formula to obtain the top-left (scalar) element of 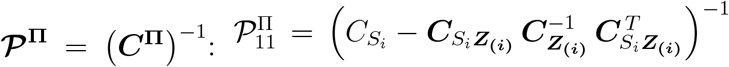. Finally, we apply the same permutation to the matrix ***P***^**Π**^. Using the fact that the permutation matrix is orthogonal and symmetric, i.e., **Π**^−1^ = **Π**^*T*^ = **Π**, we obtain: **Π *P***^**Π**^ **Π** = **Π *C***^**Π** −1^ **Π** = **Π *C***^**Π**^ **Π**^−1^ = ***C***^−1^ = ***P***. Thus, 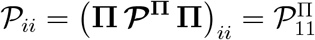.

##### Off-diagonal elements of the precision matrix

To compute the off-diagonal elements, we use a similar approach. Here, we consider the linear regression *with zero intercept* of *s*_*i*_ on ***z***_**(*ij*)**_ = (***s, k***) \ (*s*_*i*_, *s*_*j*_) (i.e., the vector of all local and contextual inputs excluding both *s*_*i*_ and *s*_*j*_). The residuals of this regression are 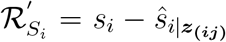, where 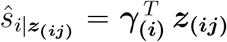 is the minimum mean squared error linear estimator of *s*_*i*_ given ***z***_**(*ij*)**_ and zero intercept. We define the scale-normalized partial cross-moment 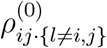 as:

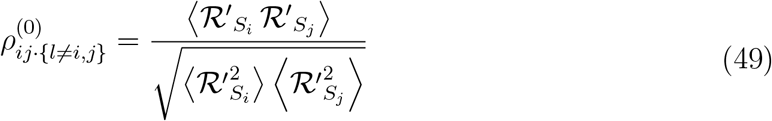

The second-moments are given by:

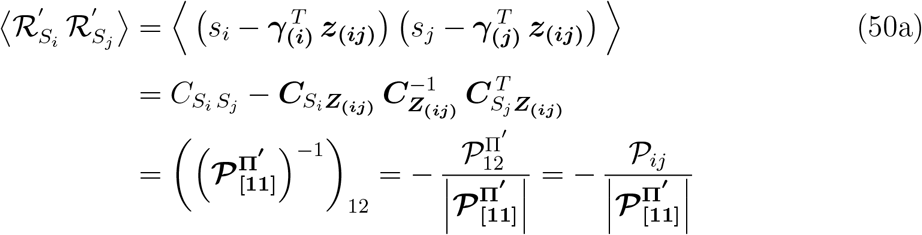

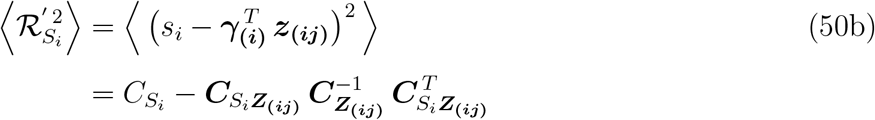

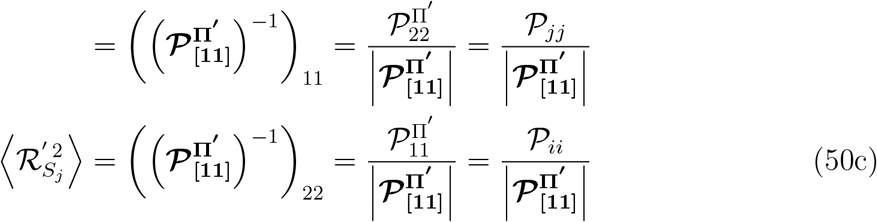

Eqs. (50) are derived analogously to Eq. (48). Here, we have 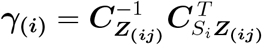, and we apply a permutation **Π**′ to the matrix ***C*** that swaps the *i*-th and first rows and columns, as well as the *j*-th and second rows and columns: ***C***^**Π**′^ = **Π**′ ***C* Π**′ . In this case, we partition ***C***^**Π**′^ so that its top-left block is 2 × 2, corresponding to the *i*-th and *j*-th elements of ***C***. We then apply the Schur complement formula to this block matrix, denoting the 2 × 2 top-left block of ***P***^**Π**′^ = (***C***^**Π**′^)^−1^ by 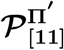. We further denote the (*k, l*) scalar element of its inverse, with *k, l* ∈ (1, 2), by 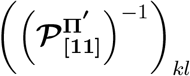. Next, we use the standard matrix inversion formula, denoting by 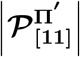 the determinant of 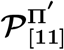 and by 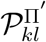 the scalar (*k, l*) element of ***P***^**Π**′^ . Finally, using the relation **Π**′ ***P***^**Π**′^ **Π**′ = ***P***, we obtain: 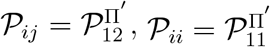, and 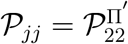.

Substituting Eqs. (50) into Eq. (49), we obtain:

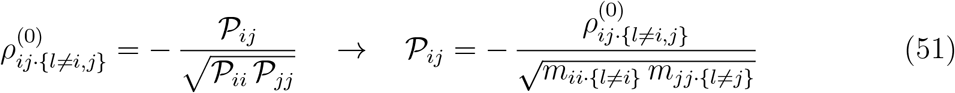

which completes the proof of the generalized form of Eqs. (47) for inverse second-moment matrices.

#### 5. Proof of formula (38b)

Formula (38b) follows directly from the definition of the top-right block of the precision matrix obtained via Schur’s inversion formula, Eq. (42), together with the equivalences established in previous sections, and the low-rank approximation 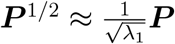:

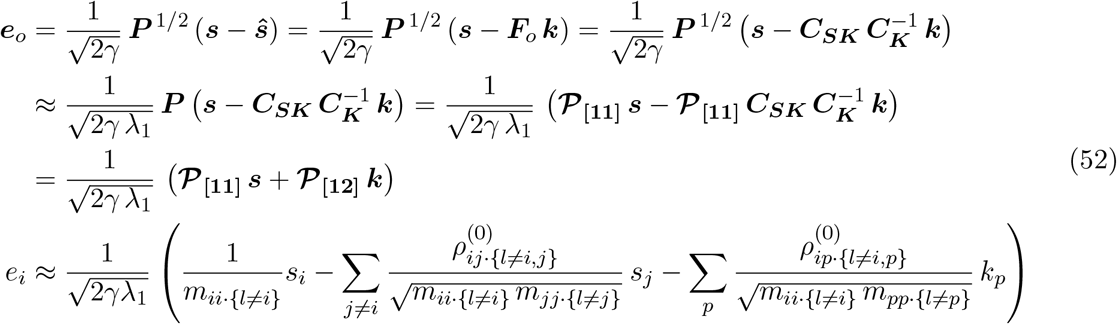

#### 6. Proof of formula (39)

Formula (39) applies when the precision matrix is diagonally dominant, that is, when ***P*** = ***D*** + ***E*** with 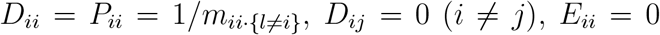, and 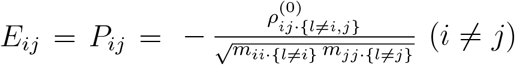, where ***E*** is small relative to ***D*** in spectral norm. We seek an approximation of the matrix square root ***P***^1*/*2^. Because ***P*** is a small perturbation of the diagonal matrix ***D***, we use a first-order Fréchet (Taylor) expansion of the matrix square root around ***D***, and write

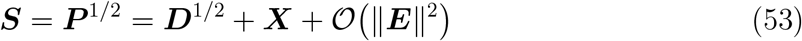

where ***X*** represents the first-order correction induced by the off-diagonal perturbation ***E***. Squaring both sides gives

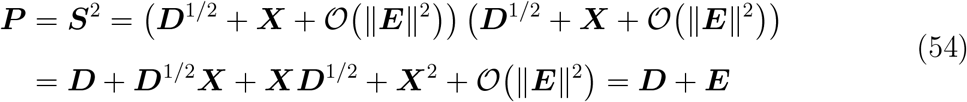

The correction ***X*** is first order in ***E***; therefore ***X***^2^ is second order and can be neglected. Matching first-order terms yields the Sylvester equation:

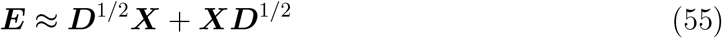

Solving for ***X***, we obtain:

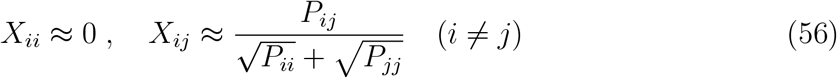

Substituting ***X*** into Eq. (53) yields the first-order approximation of the matrix square root:

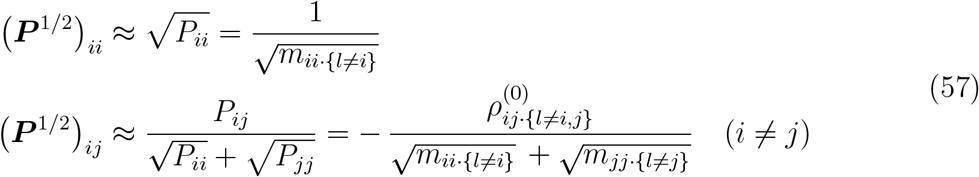

Finally, substituting formulas (57) into Eq. (37c) yields formula (39).

### D. Numerical simulations of the optimal model

#### 1. General procedure for all experiments

For all numerical simulations in Fig. 3, Fig. 4A–C, and Fig. 5, we used the following network and input parameters. We defined *N* = 100 units of type S to transmit the feedforward inputs, ***s*** = (*s*_1_, …, *s*_*N*_ ), with each input component *s*_*i*_ targeting a different E unit. Each S unit does not necessarily represent a single neuron in the brain but should be considered an effective unit transmitting a total input current to each E cell. We defined *P* = 100 units of type K to transmit the feedback (contextual) inputs, ***k*** = (*k*_1_, …, *k*_*P*_ ), with each feedback component *k*_*p*_ potentially modulating multiple E units in the effective model (Fig. 1B). We generated *T* observations (*T* = 50,000 in Fig. 3, *T* = 1,000 in Fig. 4A– C and Fig. 5) of stimuli ***s*** and contexts ***k***, corresponding to the network adaptation time window, as follows. At the times of stimulus and/or context presentation, a subset of S and/or K units was active (details of each simulation below), with amplitudes drawn from a Gaussian distribution with mean *µ* = 3 and standard deviation *σ* = 1. Each stimulus was presented at a given frequency over the adaptation time window *T* . We interpret this frequency as the relative input rate compared to that required to induce the maximum firing rate in the targeted cell, so that our maximum input frequency of 1 corresponds to inducing the highest possible firing rate in the stimulated neuron. Assuming a maximum firing rate of ∼ 500 Hz, we can estimate that input frequencies ≳ 0.02 in our model correspond to rates ≳ 10 Hz, which align with typical stimulation rates for LTP induction in the brain. In Fig. 3, we aim to simulate learning in tasks that often involve reinforcers (and thus the presence of neuromodulators, which can lower LTP thresholds); accordingly, we allow the network to adapt to lower input frequencies. White Gaussian noise *η*_*i*_ ∼ N(0, *σ*_*n*_) with *σ*_*n*_ = 0.2 was added to the input vectors ***s*** and ***k*** at all time points within the adaptation window. The network was allowed to adapt to the input statistics: the input correlation matrix 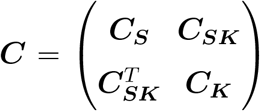 was computed over the adaptation window, and the optimal connectivity was obtained using Eqs. (4) with *γ* = 3, ensuring reasonable response amplitudes. In simulations without feedback (Fig. 4), we used the reduced model with ***k*** = **0, *F*** = **0**, and 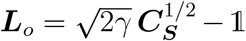. Finally, the responses of E cells, ***e***_*o*_ = (*e*_1_, …, *e*_*N*_ ), to stimuli ***s*** or conjunct stimuli and contexts (***s, k***) were obtained using Eq. (1) with the optimal connectivity. All results were robust to perturbations in parameter values.

#### 2. Audiovisual experiment

In Fig. 3A, we simulated the audiovisual experiment of [14]. Visual stimuli *V*_*a*_, *V*_*b*_, and *V*_*c*_ were simulated by activating 3 random subsets of 10% of S units, while auditory cues *A*_*a*_ and *A*_*b*_ were simulated by activating two mutually orthogonal subsets, each comprising 10% of K units. All other parameters were as described in “General procedure for all experiments”. Stimuli and contextual cues were presented at variable rates, as in [14]. Specifically, over an adaptation window of *T* = 50,000 (corresponding to 5 training days), we used the following frequencies: *f* (*A*_*a*_*V*_*a*_) = *f* (*A*_*b*_*V*_*b*_) = 0.0015 and *f* (*V*_*a*_) = *f* (*V*_*b*_) = *f* (*V*_*c*_) = 0.0003 during the first 4 days (0.8 × *T* ), and *f* (*V*_*a*_) = *f* (*V*_*b*_) = *f* (*V*_*c*_) = *f* (*A*_*b*_*V*_*a*_) = 0.0002 during the 5th day (0.2 × *T* ), such that all rates were proportional to the experimental values.

On day zero, the network was initialized to the configuration resulting from adaptation to 10 random audiovisual inputs over the window *T* . On each training day, the adaptation window was shifted by 0.2 × *T* to incorporate the stimuli presented on that day, and the network was re-optimized. After each day, we computed the responses (Resp) of E cells to *V*_*c*_, *V*_*a*_, and *A*_*a*_*V*_*a*_, as well as to *A*_*b*_*V*_*a*_ on day 5, in the optimized model.

The Response Difference Index between conditions *x* and *y* was defined as the normalized difference in E-cell responses between the two conditions, consistent with [14]: Resp. Diff. 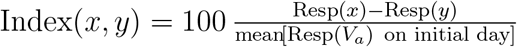. For both Resp and Resp. Diff. Index, means and standard errors (SE) were computed across tuned cells and over 50 repetitions of the experiment with different visual stimuli and auditory cues.

To confirm robustness, in Supplementary Fig. S1 we used a longer adaptation window (*T* = 500,000), correspondingly 10-fold lower frequencies, and 2 random (rather than orthogonal) subsets of 10% of K units to simulate each auditory cue.

#### 3. Audiomotor experiments

In Fig. 3BC, we simulated the audiomotor experiments of [35] and [37]. Six sounds (1 reafferent, referred to as “expected”, *S*_expected_, and 5 deviant, *S*_deviant_) were simulated by activating 6 random subsets of 10% of S units, while motor feedback (*M* ), representing the efference copy signal during running or lever pressing, was simulated by activating a random subset of 10% of K units. All other parameters were as described in “General procedure for all experiments”.

To simulate audiomotor training, we presented each sound in the presence or absence of motor feedback at the following frequencies over an adaptation window of *T* = 50,000: *f* (*MS*_expected_) = 0.0025 (expected sound during movement), *f* (*MS*_deviant_) = 0.00002 (each of the five deviant sounds during movement), and *f* (*S*_deviant_) = 0.00002 (each of the five deviant sounds at rest). These rates were chosen to be proportional to those used across the three experimental phases on each training day in [35]. The experiment in [37] was similar and we therefore used the same protocol.

We allowed the network to adapt to these input statistics and then computed the responses of E cells to the expected and deviant sounds during movement and at rest (i.e., in the presence or absence of motor feedback, respectively) in the optimized model. In Fig. 3B, we plotted the responses of E cells tuned to the expected sound and to one deviant sound for each of 50 repetitions of the numerical experiment.

In Fig. 3C, we computed the responses of I cells (using the procedure described in the section below) to the expected sound, a deviant sound, and the motor feedback alone, under two conditions: “Trained”, corresponding to the protocol described above, and “Naive”. For the latter, we simulated adaptation to a period of movement without sound (*f* (*M* ) = 0.0025 over the adaptation window). Fig. 3C shows the responses of *Q* = 25 I cells (1/4 of the number *N* of E cells, consistent with typical I/E ratios in cortex [94]) for each of 50 repetitions. For each condition, we computed regression slopes and Cohen’s *f*^2^, together with their confidence intervals (CI), using a bootstrap analysis with 1,000 samples.

In Supplementary Fig. S1, we used a longer adaptation window (*T* = 500,000) and correspondingly 10-fold lower frequencies.

#### 4. Visual–olfactory experiment

In Fig. 3D, we simulated the visual–olfactory experiment of [15]. An odor (*O*_*a*_) was simulated by activating a random subset of 10% of S units, and 2 visual contexts (*V*_*a*_ and *V*_*b*_) were simulated by activating two mutually orthogonal subsets, each comprising 10% of K units. All other parameters were as described in “General procedure for all experiments”.

We presented the odor input in the first visual context at the following frequency over an adaptation window of *T* = 50,000: *f* (*O*_*a*_*V*_*a*_) = 0.0025.

We allowed the network to adapt to these input statistics and, using the optimized model, we computed the responses of E cells in three conditions: *O*_*a*_*V*_*a*_ (odor in the familiar context), *V*_*a*_ (familiar context alone), and *V*_*b*_ (novel context alone). We then used these E-cell responses to estimate the responses of *Q* = 1,000 I cells in each condition, using the approach described in the next section. Here, we considered a number of I cells 10 times greater than E cells, in line with the substantially larger number of granule cells (I) compared with mitral and tufted cells (E) in the olfactory bulb [150]. All negative I responses were set to zero, consistent with the fact that the Zif268 early gene used in [15] detects only positive changes in neural activity.

Fig. 3D (right) shows scatter plots of I cell responses in the different conditions across 50 repetitions of the numerical experiment. We then computed the average response over I cells for each repetition. Fig. 3D (left) shows the mean ± SD of this average response across repetitions in each condition. We further computed response overlap as the cosine similarity (in %) between I cell responses in the (*O*_*a*_*V*_*a*_, *V*_*a*_) and (*V*_*a*_, *V*_*b*_) conditions. Fig. 3D (middle) shows the mean ± SD of these overlaps across repetitions.

In Supplementary Fig. S1, we used 2 non-orthogonal subsets of active K units (each comprising 10% of all K units) to simulate each visual context. Of these active units, 20% were assigned identical responses in both contexts, while the remaining components were chosen to be orthogonal. This design was intended to reflect visual contexts with shared features as well as distinct features, as in [15].

#### 5. Estimation of I-cell responses via constrained connectivity inference

##### Inference of synaptic connectivity under Dale’s law

To compute I-cell responses in each condition, the first step was to estimate the synaptic weights between I neurons and E and K cells that satisfy the sign constraint imposed by Dale’s law and generate effective connectivity matrices ***L*** and ***F***, Eqs. (10), closely approximating the optimal matrices ***L***_*o*_ and ***F***_*o*_, Eqs. (20). For simplicity, we considered circuits without direct I-I, E-K, and cross E-E connections, while retaining all other connections shown in Fig. 1A. We thus searched for approximate solutions to:

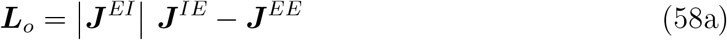

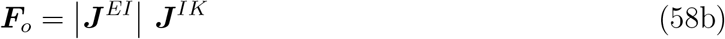

subject to:

- (Dale sign constraint) 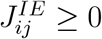 and 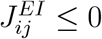 for all *i, j*; 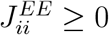 for all *i*;
- (symmetry) ***J***^*EI*^ = −(***J***^*IE*^)^*T*^ ;
- (no E–E cross-coupling) ***J***^*EE*^ is diagonal.

##### Nonnegative matrix factorization via projected gradient descent

To estimate ***x*** =|***J***^*EI*^|, we minimized the squared Frobenius norm of the residuals of Eq. (58a),

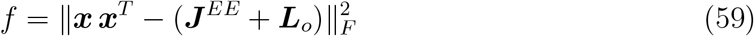

subject to *x*_*ij*_ ≥ 0 for all *i, j*. Optimization was performed using projected gradient descent (PGD) with elementwise projection onto the nonnegative orthant after each update and learning rate *α* = 10^−4^. Iterations were terminated when the relative change in Frobenius norm, 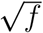, fell below 10^−6^. Convergence to a stationary solution was achieved in all simulations.

##### Initialization via spectral factorization

To initialize PGD, we proceeded as follows.

First, we selected ***J***^*EE*^ so that the target matrix ***J***^*EE*^ + ***L***_*o*_ is positive definite, ensuring that the factorization problem is well-posed. Using Gershgorin’s circle theorem, we set

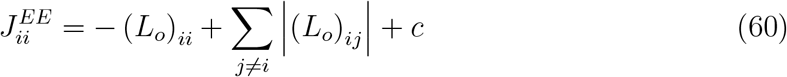

with 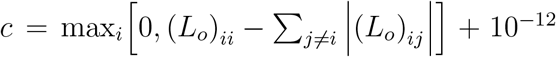, and 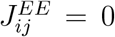 for *i*≠*j*. This choice makes ***J***^*EE*^ + ***L***_*o*_ strictly diagonally dominant with positive diagonal entries (and hence positive definite, given matrix symmetry), and ensures 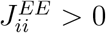 for all *i*.

Second, we analytically minimized *f* without enforcing the sign constraint on ***x***, as follows.

- When *Q* ≥ *N* (visual–olfactory experiment), one exact solution (with *f* = 0) is

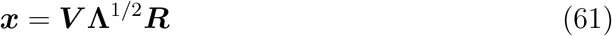

where ***V*** ∈ ℝ^*N* ×*N*^ contains the eigenvectors of ***J***^*EE*^ + ***L***_*o*_, **Λ** ∈ ℝ^*N* ×*N*^ is the corresponding diagonal matrix of eigenvalues, and ***R*** ∈ ℝ^*N* ×*Q*^ is a random matrix with orthonormal rows.
- When *Q < N* (audiomotor experiments), an exact solution generally does not exist. We therefore used the truncated approximation

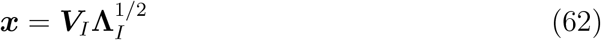

where ***V***_*I*_ = ***V***_[:,1:*Q*]_ ∈ ℝ^*N* ×*Q*^ and **Λ**_*I*_ = **Λ**_[1:*Q*,1:*Q*]_ ∈ ℝ^*Q*×*Q*^, corresponding to the leading *Q* eigenmodes of ***J***^*EE*^ + ***L***_*o*_.

In both cases, we then set any negative elements of ***x*** to zero and used the resulting matrix to initialize PGD.

##### Reconstruction of Dale-compliant connectivity

After convergence, PGD yields a Dale-compliant solution,|***J***^*EI*, Dale^ | = ***x*** and ***J***^*IE*, Dale^ = ***x***^*T*^ . We then computed the corresponding diagonal E-E weights from Eq. (58a):

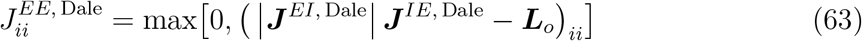

Finally, we obtained the K-to-I connectivity from Eq. (58b) as:

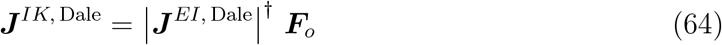

where ^†^ denotes the Moore-Penrose pseudoinverse.

##### Computation of I-cell responses from inferred connectivity

We reconstructed the circuit effective connectivity from the inferred synaptic weights:

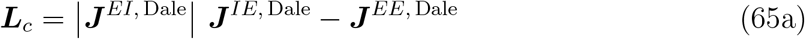

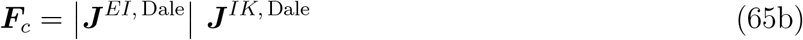

Supplementary Fig. S2 shows that, although exact identity cannot be achieved due to the imposed constraints, ***L***_*c*_ and ***L***_*o*_, as well as ***F***_*c*_ and ***F***_*o*_, are highly correlated. As a result, E-cell responses computed from the reconstructed (constrained) connectivity closely match those obtained from the normative model.

We then estimated the I-cell responses (***i***) using the circuit equation:

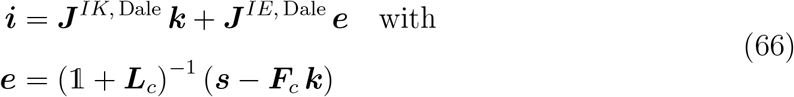

where ***s*** and ***k*** are determined by the experimental conditions. Specifically, in Fig. 3C, we set ***k*** = *M*, i.e., the motor feedback, and ***s*** = **0** (y-axis, all conditions); ***k*** = **0** and ***s*** = *S*_expected_ (x-axis, first “Trained” condition and “Naive”; note that for the “Naive” condition the choice of sound is irrelevant); ***k*** = **0** and ***s*** = *S*_deviant_ (x-axis, second “Trained” condition). In Fig. 3D, we used ***s*** = *O*_*a*_ and ***k*** = *V*_*a*_ (*O*_*a*_*V*_*a*_ condition), ***s*** = **0** and ***k*** = *V*_*a*_ (*V*_*a*_ condition), and ***s*** = **0** and ***k*** = *V*_*b*_ (*V*_*b*_ condition).

#### 6. Neural habituation

In Fig. 4A, we simulated a stimulus ***s*** by activating 10% of the S units at a constant rate (input frequency on the x-axis) over the adaptation window *T*, with all other parameters as described in “General procedure for all experiments”. We then optimized the model and computed the responses of E cells to the stimulus. We iterated the experiment across different input frequencies and 10,000 random stimuli, calculating the mean and interquartile range (IQR) of responses across cells and stimuli. In Fig. 5A, we simulated a context by activating 10% of the K units concurrently with the S units. The K units were activated at a relative context-to-stimulus rate of 0.2 (i.e., once every 5 feedforward stimulations ***s***, representing a ‘weak context’ association) or at a relative rate of 1 (i.e., at every feedforward stimulation, representing a ‘strong context’ association). We optimized the model based on these statistics and computed the E-cell responses to the same stimulus + context. Results were averaged over 10,000 iterations of the experiment with different random stimuli and contexts.

#### 7. Shift in neural tuning

In Fig. 4B, model connectivity was optimized based on 50 independent stimuli, each presented at a constant rate of 0.02 over the adaptation window. The responses of E cells to these 50 familiar inputs and 50 novel inputs were then computed in the optimized model. Each E cell was directly activated by 5 random inputs from the familiar set and 5 random inputs from the novel set – i.e., 10% of all stimuli – corresponding to the activation, on average, of 10% of the S units for each stimulus ***s***. For each E cell, the 5 familiar inputs were ranked in ascending order and the 5 novel inputs were ranked in descending order. The mean and interquartile range (IQR) of inputs and responses were then computed across 100 cells and 10,000 repetitions of the experiment with varying stimuli. In Fig. 5B, we simulated a distinct context ***k*** for each stimulus ***s*** by activating 1 K unit concurrently with the S units. A different K unit was activated for each of the 100 stimuli. For the 50 familiar stimuli, the K unit was activated at a relative context-to-stimulus rate of 0.2 (‘weak context’) or 1 (‘strong context’) during the adaptation window, and the connectivity was optimized based on these statistics. Responses to each stimulus were computed in its corresponding context.

#### 8. Pattern separation and inverse effectiveness

In Fig. 4C and Fig. 5C–G, we generated two stimuli, ***s***^(1)^ = (*A, B*) and ***s***^(2)^ = (*A, C*), each consisting of a shared component (*A*) and an independent component (*B* or *C*, respectively). Each of the three components corresponded to a distinct, non-overlapping set of 10% active S units. The two stimuli were repeatedly presented in an alternating sequence at a constant frequency during the adaptation window. The experiment was then iterated with varying input frequencies. In Fig. 4C, the model connectivity was optimized for each input frequency (x-axis) and the cosine distance between the E-cell responses to ***s***^(1)^ and ***s***^(2)^ was calculated (y-axis). In Fig. 5, we simulated each context by activating 10% of the K units concurrently with the associated stimulus, at a relative context-to-stimulus rate of 0.2 (‘weak context’) or 1 (‘strong context’) during the adaptation window. The cosine distance between the contexts associated with the two stimuli ranged from 0 (identical contexts, Fig. 5C) to 1 (orthogonal contexts, Fig. 5DE), spanning intermediate values (Fig. 5FG). The connectivity was optimized based on the statistics of stimuli and contexts, for each input frequency. The cosine distance between E-cell responses to ***s***^(1)^ and ***s***^(2)^ (in their respective contexts) was computed (Fig. 5C–F), and the mean E-cell response to each input component (*A, B*, and *C*) was plotted for a stimulus frequency of 0.02 (Fig. 5CDE). The mean and IQR of responses and cosine distances were generally evaluated over 10,000 repetitions with different stimuli and contexts. For the full range of context scenarios (Fig. 5F), we used 5,000 repetitions.

In Fig. 5D, the disinhibition of the component absent from the input, compared to the case without context, occurs because the suppression of *A* disinhibits *B* and *C* via the lateral connectivity ***L***_*o*_. In Fig. 5E, the amplification of the component specific to each input, relative to the case without context, arises because both contexts predict *A*, which in turn enhances the response to the non-predicted component through ***L***_*o*_. In Fig. 5G, the discriminability of independent representations (x-axis) quantifies the overall discriminability of stimuli and contexts from disconnected neural representations (i.e., without the feedback interaction). In familiar contexts, this is defined as the *sum* of the discriminability between E-cell responses to ***s***^(1)^ and ***s***^(2)^ (without feedback) and the discriminability of the context representations: 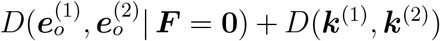, where *D* denotes the cosine distance. In swapped contexts, it is defined as the *difference* between the two discriminabilities: 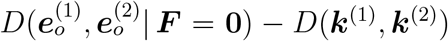. This difference arises because, from the perspective of a downstream decoder, distinct contexts reduce sensory ambiguity when the information is congruent across modalities, i.e., when stimulus-context associations are familiar. In contrast, distinct contexts increase sensory ambiguity when the information is incongruent across modalities, i.e., when stimulus-context associations are swapped. The discriminability gain from contextual feedback (y-axis) is defined as the change in overall discriminability resulting from the feedback connectivity. Since only the ***e***_*o*_ representations are affected, this gain is the difference in discriminability between 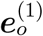 and 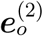 with and without feedback: 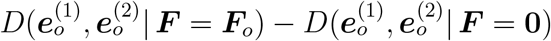.

## ACKNOWLEDGMENTS

The author thanks Martha Bagnall, Ilya Monosov, ShiNung Ching, Ryan McGee, and Baranidharan Raman for fruitful discussions. This research was supported by the Alfred P. Sloan Foundation (Grant No. FG-2024-22163).

## APPENDIX: Generalization to a variance-based cost function

The equivalence between context-conditioned efficient coding and predictive coding generalizes to the case in which the cost term in multimodal Efficiency penalizes response fluctuations rather than response magnitude:

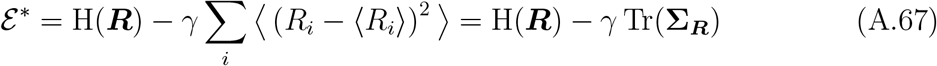

In this case, Eqs. (37) become:

Multimodal Efficient Coding ≡ Multimodal Predictive Coding

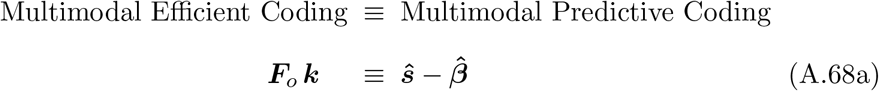

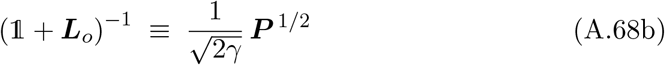

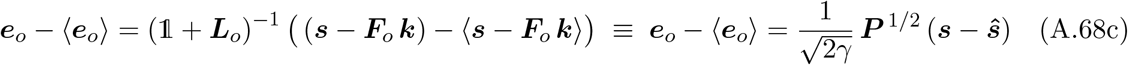

with the following changes in interpretation:

1. the optimal connectivity, ***L***_*o*_ and ***F***_*o*_, is given by Eqs. (20) with the second-moment matrices replaced by the corresponding covariance matrices;
2. 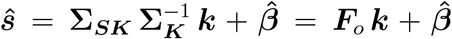 is the vector of optimal linear predictions computed *with intercepts* 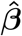; the intercepts are also automatically optimized, i.e., 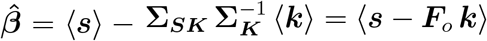;
3. the precision matrix is interpreted in the classical sense, that is, ***P*** = ***P***_[11]_ = (**Σ**^−1^)_[11]_. The partial variances and partial correlations in the precision matrix, Eqs. (47), are the variances and Pearson’s correlations of the residuals of the linear regressions *with intercepts*. For example, each linear regression of *s*_*i*_ on ***z***_**(*i*)**_ = (***s, k***) \ *s*_*i*_ includes an intercept 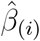, leading to residuals 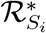 where the inputs are centered:

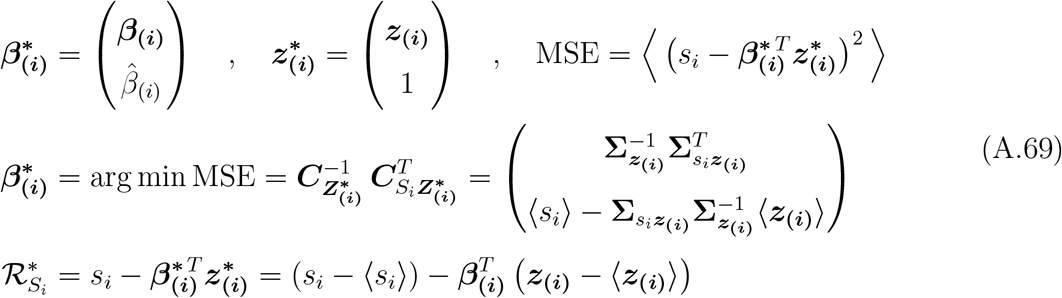

(The other residuals are defined similarly).

Eq. (A.68c) highlights that, in this case, mixtures of prediction errors are encoded in fluctuations of E-neuron activity, rather than in their raw activity levels.

This encoding has similar properties to Eq. (37c), with some nuanced differences.

First, the representation is whitened with respect to the covariance matrix **Σ**_***ε***_ of the prediction errors ***ε*** = ***s*** − ***ŝ***:

Proof:

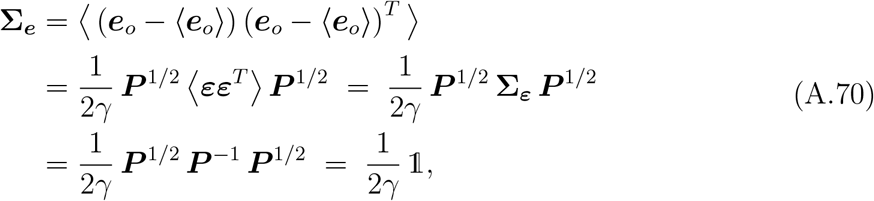

where we used ⟨***ε***⟩ = **0** and:

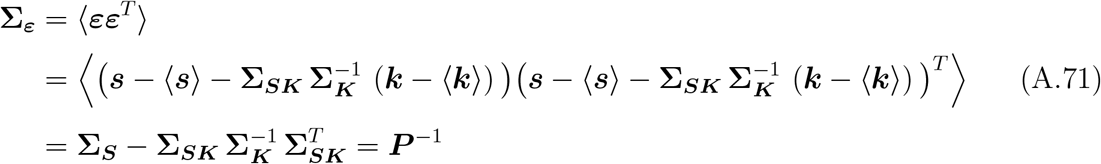

Second, the Euclidean norm of the population response fluctuations, ∥***e***_*o*_ − ⟨***e***_*o*_⟩∥, is proportional to the Mahalanobis distance *d*_*M*_ (***ε***) of the prediction error vector ***ε***, that is, the distance of prediction errors from the center of their distribution in units of standard deviations. This provides a direct probabilistic interpretation of the squared norm of response fluctuations: under our Gaussian assumption on the input distribution, prediction errors are also normally distributed, ***ε*** ∼ N (**0, Σ**_***ε***_), and the squared magnitude of response fluctuations corresponds to the negative log-likelihood of the prediction errors (up to a rescaling factor and an additive constant). Thus, it quantifies how statistically surprising the prediction errors are.

Proof:

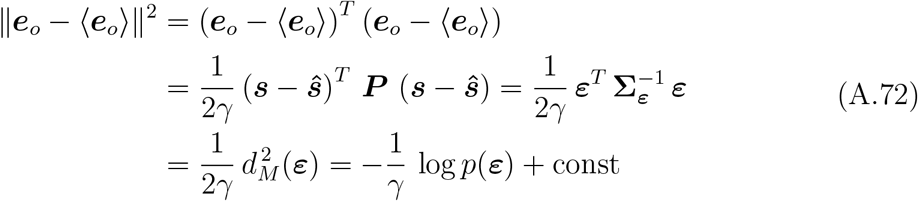

Both properties generalize to the entire class of degenerate solutions ***e***^′^ = ***Q e***_*o*_, since orthogonal transforms preserve norms and whitening.

