## Supplemental_Material for "A unified theory of context-conditioned efficient and predictive coding"

Gaia Tavoni

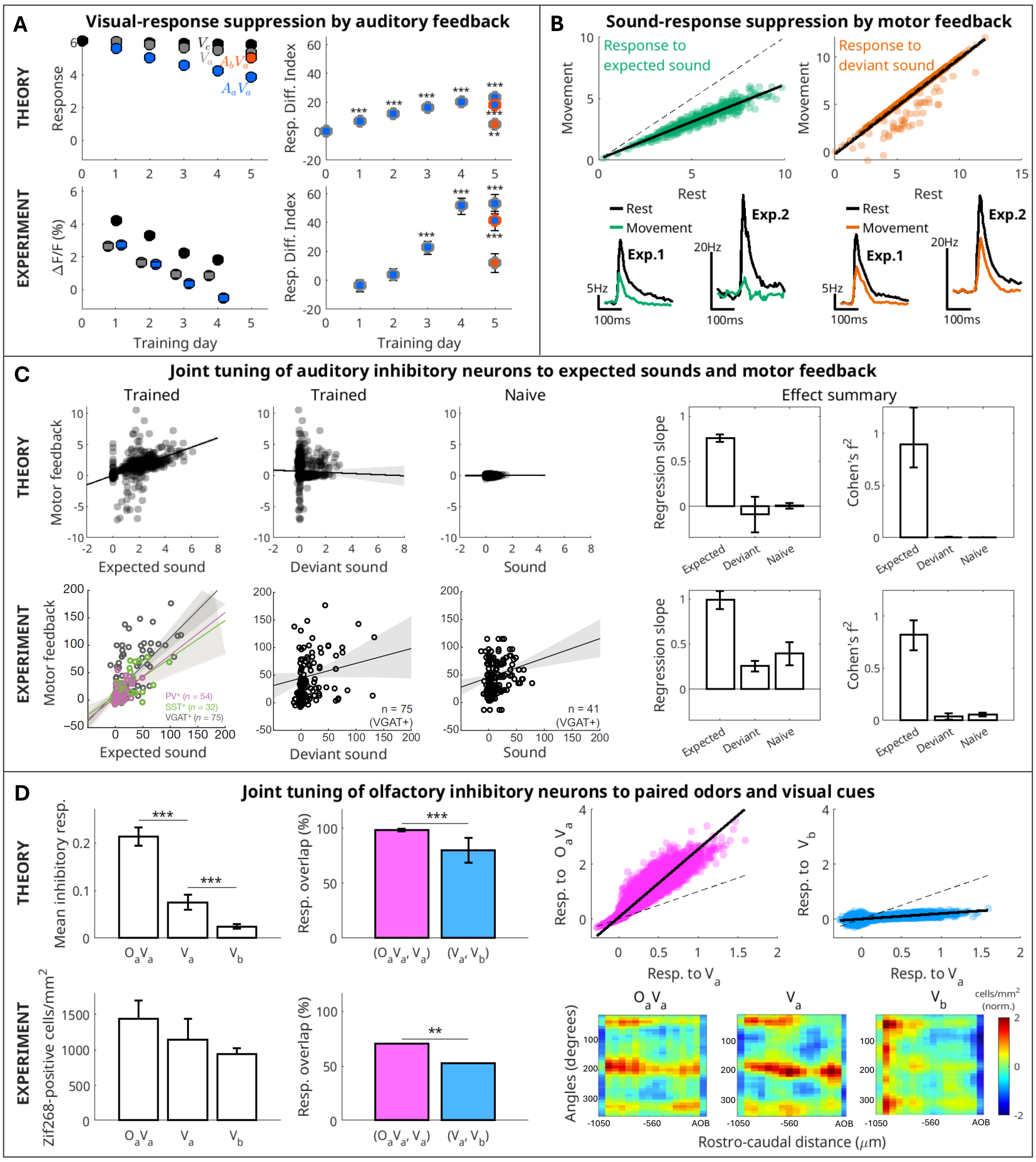


Figure S1: **Results in Fig. 3 are robust to changes in stimulus frequency and cross-modal context overlap**. (**A–C**) Same audiovisual and audiomotor experiments as in Fig. 3A and Fig. 3B–C, respectively, but with a longer adaptation window in the numerical simulations ($T = 500,000$) and correspondingly 10-fold lower input and feedback frequencies. Additionally, in (A), two random (rather than orthogonal) subsets of 10% of K units are used to encode the two auditory cues ($A_a$ and $A_b$). (A–B) At lower frequencies, E-cell responses are less strongly suppressed by cross-modal context, as expected. (A–C) Predicted effects (top) remain consistent with the experimental data (bottom). (**D**) Same visual–olfactory experiment as in Fig. 3D, but using two non-orthogonal subsets of active K units (each comprising 10% of the population) to encode each visual context. Of these active units, 20% have identical responses across contexts, while the remaining components are orthogonal. This manipulation increases the overlap in I-cell responses to $V_a$ and $V_b$ (top), in agreement with the experimental data (bottom).

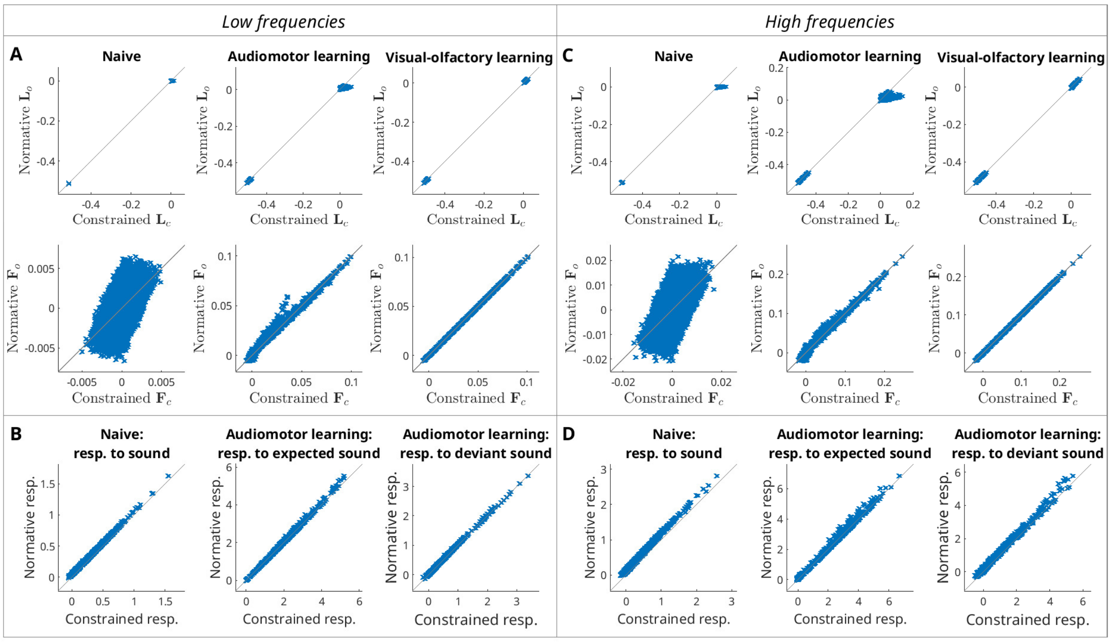


Figure S2: **Connectivity and E-cell responses inferred under Dale's law closely match the normative solution.** Dale-compliant synaptic weights and the corresponding constrained effective connectivity ($\boldsymbol{L}_c$ and $\boldsymbol{F}_c$) are inferred using a hybrid analytical-numerical approach to approximate the normative solution (Methods). (**A, C**) The inferred connectivity matrices, $\boldsymbol{L}_c$ and $\boldsymbol{F}_c$, are highly correlated with their normative counterparts, $\boldsymbol{L}_o$ and $\boldsymbol{F}_o$. Results are shown for the audiomotor and visual–olfactory experiments of Fig. 3CD. (**B, D**) E-cell responses are nearly identical whether computed from the constrained connectivity or directly from the normative solution. Results are shown for the audiomotor experiment of Fig. 3C. Results are obtained using (A, B) lower input and feedback frequencies, as in Supplementary Fig. S1, and (C, D) higher frequencies, as in Fig. 3CD (Methods).
